# A transition-prone brain state precedes spontaneous behavioral switching

**DOI:** 10.64898/2026.03.06.709845

**Authors:** Paulina Wanken, Bradley Jay Edelman, Leafy Behera, Jose Maria Martinez de Paz, Patrick T. McCarthy, Emilie Macé

**Author notes:** Correspondence to: Emilie Macé. Current addresses: Leafy Behera, Department of Biomedicine, University of Basel, Basel, CH, Patrick T. McCarthy, Centre for Neural Circuits and Behaviour, University of Oxford, Oxford, UK.

## Abstract

Animals exhibit behavior in the absence of external stimuli or explicit tasks. Is the initiation of such spontaneous behavior shaped by internal brain states in a predictable manner? If so, does it engage specific brain circuits independent of behavioral form? Here, we studied the initiation of uninstructed behaviors of head-fixed mice in two contexts: a virtual burrow and a running wheel. Across both contexts, mice spent most of the time in quiet wakefulness and spontaneously initiated bouts of egress (exiting the burrow), running, or grooming. We employed functional ultrasound imaging (fUS) to record whole-brain activity and to identify whether the initiation of spontaneous behavior could be predicted from hemodynamic signals. We first identified distinct hemodynamic patterns associated with each behavior and subsequently performed time-resolved decoding to predict behavioral transitions from fUS data. We found that whole-brain hemodynamic signals could decode spontaneous egress and running around 10 seconds before their onset, a timescale that cannot be accounted for by preceding behavioral changes alone. Furthermore, we found a network of regions, including the medial septum (MS), that decreased their signal several seconds before the onset of egress and running. Mimicking this decrease by inhibiting neurons in the MS via optogenetics increased the probability of egress, running, and grooming. Through this unbiased approach, our work sheds light on a whole-brain transition-prone state that precedes uninstructed behavior transitions.

## Introduction

Being able to switch behaviors is fundamental to all animals’ survival, allowing them to meet physiological needs and adapt to changing environments. Transitioning between behavior states, such as from exploration to exploitation^1^, or between active and inactive states^2^, requires the brain to integrate diverse streams of information, including sensory cues, memory, and internal state. This points to the involvement of distributed brain areas, yet the brain-wide mechanisms that orchestrate spontaneous behavior transitions remain poorly understood. Do they arise stochastically, or are they the result of a gradual change in internal state? Are specific brain regions involved in promoting spontaneous transitions in behavior? Studying spontaneous, uninstructed behaviors that are not coupled to specific learned tasks offers the opportunity to address these questions.

In humans and non-human primates, neural recordings that predict spontaneous actions suggest that the brain undergoes coordinated changes before behavior onset. Seminal electrophysiological studies identified a so-called readiness potential in the pre-and supplementary motor areas hundreds of milliseconds before movement ^3,4^. More recent work examining whole-brain activity in humans using fMRI, showed that the choice between pressing a left or right button can be decoded in distinct cortical regions seconds in advance^5^. These studies have greatly advanced our understanding of spontaneous behavior initiation, converging on the idea that behavioral transitions are predictable before an action occurs. Other fMRI studies in primates have shown that specific large-scale networks, such as the salience network, may play a crucial role in detecting salient information, reorienting attention, and initiating shifts in behavioral state^6–8^. However, with limited ability to manipulate such neural circuits, it remains challenging to test the causal role of associated regions in behavior transitions.

In rodents, research on behavioral transitions has largely focused on individual regions, such as the prefrontal cortex^9–13^ and striatum^9,14,15^. Furthermore, neuromodulatory systems - including the noradrenergic^16–21^, serotonergic^22–24^, and cholinergic system^25–27^ - have been implicated in behavioral switching. While these studies have provided valuable insights into region-and task-specific mechanisms, they often lack a brain-wide perspective and thus may fail to capture broad activity patterns supporting uninstructed behavioral transitions. Widefield cortical imaging in mice has shown that upcoming behavioral switches can be predicted from cortical activity seconds in advance^28^, but the role of subcortical regions remains unclear. Moreover, fMRI in rodents has been mostly restricted to anesthetized states or constrained behaviors^29,30^, leaving spontaneous behavior relatively unexplored. A brain-wide perspective would capture distributed activity patterns that support uninstructed behavioral transitions, while benefiting from circuit manipulation tools available in mice.

Here, we address this gap by investigating the neural basis of how mice transition from quiescence to movement in the absence of external triggers at the brain-wide level. To this end, we employed functional ultrasound imaging (fUS) to record whole-brain information while mice engaged in spontaneous behaviors in two different head-fixed contexts. fUS measures a hemodynamic signal that is correlated with the underlying neuronal activity^31,32^. We observed spontaneous initiations of egress and grooming in a virtual burrow, and of running on a wheel, and identified whole-brain correlates associated with each of these behaviors. Next, we applied machine learning to the fUS data and found that behavioral transitions can be decoded from brain-wide activity around 10 seconds in advance. Moreover, we found that a distributed network of regions exhibited a decrease in hemodynamic signal seconds before behavior onset. Finally, we performed optogenetic inhibition of the Medial Septum (MS), a candidate switching region identified through fUS, and found that it facilitates behavior transitions.

## Results

### Head-fixed mice spontaneously initiate stereotyped behaviors

To examine spontaneous behavioral transitions in the absence of external stimuli, we first used the Virtual Burrow Assay (VBA)^33^ while recording whole-brain activity via fUS. This assay aims to model the innate behavior of mice in a burrow under head-restrained conditions. In the VBA, head-fixed mice are placed inside an air-lifted tube that they can voluntarily exit to explore a small object coupled to the burrow (Fig. 1a). We refer to this behavioral state as egress (Fig. 1a). Multiple behavioral variables were recorded in this context: the position of the burrow and the force exerted on it, as well as pupil size and face motion^34^ through additional cameras.

**Fig. 1.**
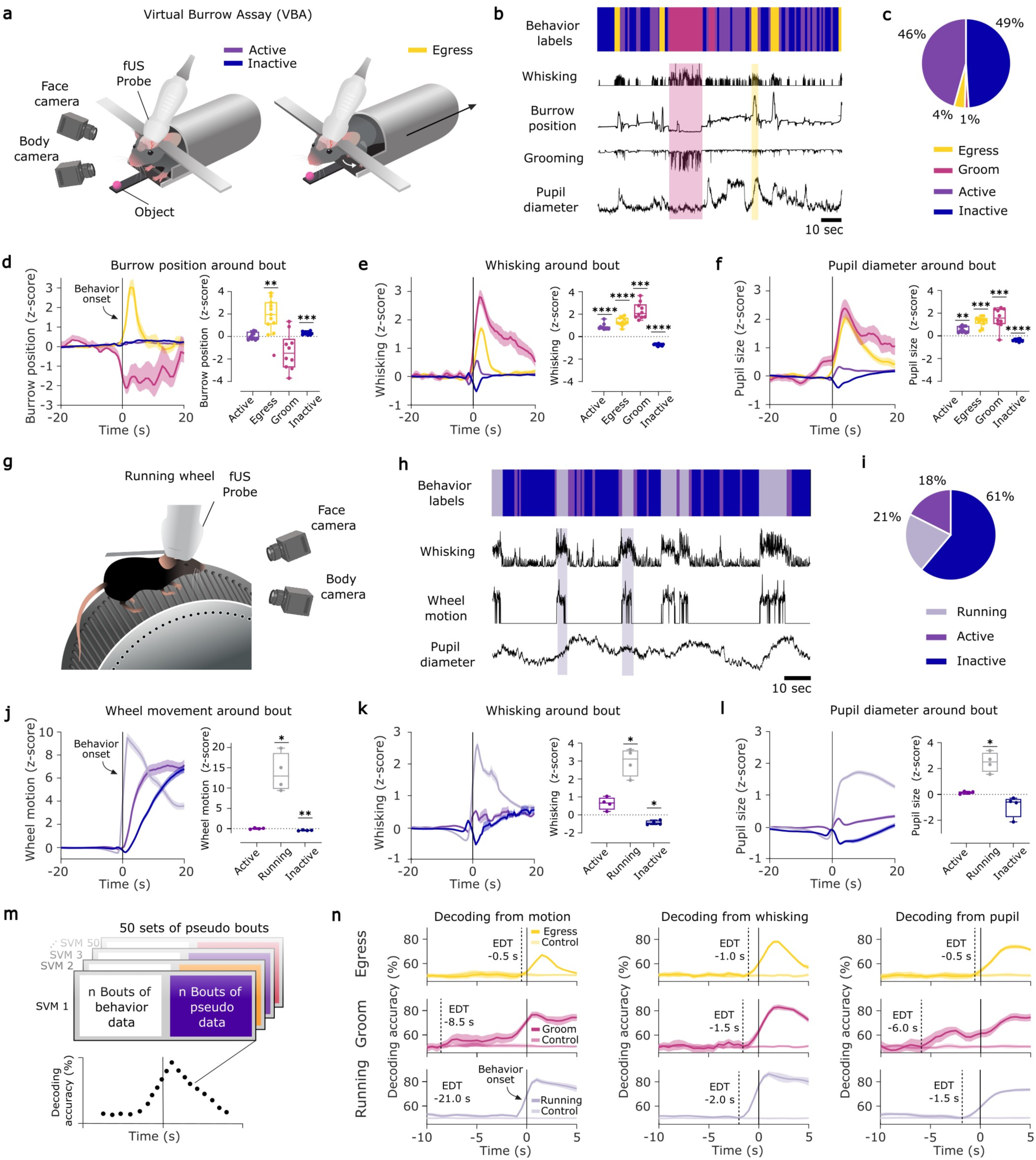
Behavioral states in the Virtual Burrow Assay (VBA) and on the running wheel. **a**, Schematic of VBA setup used for observing spontaneous behavior and functional ultrasound imaging. Mouse in active or inactive (left) and in egress state (right). **b**, Example behavioral readouts and corresponding color-coded behavioral states on top. **c**, Proportion of behavioral states in the VBA across all sessions (n = 55, egress = 1.37 %, groom = 3.90 %, active = 45.69 %, inactive = 49.01 %). **d**, VBA motion around onset of egress (yellow line, mean ± s.e.m. across n = 11 animals for all states from **c-f**), grooming (pink line), active (purple line), and inactive (blue line) (left) and quantification (right). Mixed-effects model analysis with Tukey’s multiple comparisons test (**d-f** and **j-l**), comparing VBA motion during behavior to baseline, active vs. baseline (p = 0.6634), egress vs. baseline (p = 0.0029), groom vs. baseline (p = 0.1695), inactive vs. baseline (p =.0.0003). **e**, Whisking around onset of states (left) and quantification (right). Comparing whisking during behavior to baseline, active vs. baseline (p < 0.0001), egress vs. baseline (p < 0.0001), groom vs. baseline (p = 0.0005), inactive vs. baseline (p < 0.0001). **f**, Pupil diameter around onset states (left) and quantification (right). Comparing pupil size during behavior to baseline, active vs. baseline (p = 0.0029), egress vs. baseline (p = 0.0001), groom vs. baseline (p = 0.0009), inactive vs. baseline (p < 0.0001). **g-i,** The same as (**a-c**), except for running wheel (n = 35, running = 17.54 %, active = 21.33%, inactive = 61.12%). **j**, Wheel motion around onset of running (gray line, mean ± s.e.m. across n = 4 animals), active (purple line), and inactive (blue line) (left) and quantification (right). Comparing wheel motion during behavior to baseline, active vs. baseline (p = 0.9986), running vs. baseline (p = 0.0455), inactive vs. baseline (p = 0.0092). **k**, Whisking around onset of states, (left) and quantification (right). Comparing whisking during behavior to baseline, active vs. baseline (0.1884), running vs. baseline (p = 0.0211), inactive vs. baseline (p = 0.0132) **l**, Pupil diameter around onset states (left) and quantification (right). Comparing pupil size during behavior to baseline, active vs. baseline (p = 0.4479), running vs. baseline (p = 0.0378), inactive vs. baseline (p = 0.3718). **m**, Schematic of behavior decoding using classification SVMs. **n**, Average decoding accuracies resulting from SVM decoding from burrow position (equivalent to motion in VBA) (left), whisking (middle) and pupil size (right) around egress (yellow line, mean ± std across 50 SVM runs) and grooming (pink line) and wheel motion (left), whisking (middle) and pupil size (right) for running (gray line). Lighter versions of yellow, pink and gray represent 50 control SVM runs. Dashed lines indicate the earliest decoding time (EDT), defined as the first time at which the mean behavior decoding accuracy falls within one standard deviation of the control decoding accuracy (pseudo-bout vs. pseudo-bout). * p < 0.05, ** p < 0.01, *** p < 0.001, **** p < 0.0001.

First, we classified the behavior of 11 mice across 55 sessions into four distinct states. The mice exhibited two infrequent, long lasting, stereotyped behaviors: egress (1% of the time, 3.9 s mean duration) and groom (4%, 9.3 s) (Fig. 1b, Extended Data Fig. 1b). Egress was detected using a threshold on the speed of the burrow in the VBA, while grooming was identified using a video correlation-based approach within a snout region-of-interest. The rest of the time (95%) (Fig. 1c), the mice were in a less mobile, quiet wakefulness state, which could be decomposed into relatively short active (46%, 2.7 s) and inactive states (49%, 2.8 s) based on whisking activity (Fig. 1b, Extended Data Fig. 1b) extracted via a motion singular value decomposition (SVD) based readout (from FaceMap)^28^ of the facial video. We did not observe an effect of cumulative session time on the occurrence of active and inactive states. Similarly, egress and groom bouts occurred in an apparently spontaneous, random manner throughout a session (Extended Data Fig. 1a), providing a useful framework to investigate the uninstructed initiation of behaviors.

To examine the difference between all states, we quantified burrow position, whisking activity and pupil size during each of the states across mice. As expected, based on our classification criteria, burrow position changed sharply at the onset of egress. In contrast, burrow position remained relatively stable during active and inactive states (Fig. 1d). Pupil size and whisking activity showed similar dynamics, with an increase during the active state and a decrease during the inactive state. Additionally, we observed a strong increase in both whisking activity and pupil diameter following the onset of egress and grooming (Fig. 1e-f).

To extend our investigation to other behavioral transitions, we included an independent, complementary cohort on a running wheel. In this context, we recorded whole-brain fUS from head-fixed mice (n = 4) that were able to run spontaneously (Fig. 1g). As in the VBA setup (Fig. 1a-c) we classified quiet wakefulness into active and inactive states based on whisking, and added a third state, running, defined by thresholding the motion of the wheel (Fig. 1g-h). These states again occurred randomly distributed throughout a session (Extended Data Fig. 1c). As in the VBA, mice spent most of their time on the wheel either in the inactive (61%) or active (18%) state (Fig. 1i), while they ran 21% of the time. Notably, in this cohort mice did not exhibit grooming.

Running bouts were on average the longest in this context, with a mean duration of 8.8 seconds (Extended Data Fig. 1d). As expected, based on its definition, wheel motion increased at the onset of running (Fig. 1j), and was accompanied by an increase in both whisking (Fig. 1k) and pupil diameter (Fig. 1l) as previously reported in literature^35,36^. The average duration of active bouts (2.5 s mean duration) was slightly shorter than that observed in the VBA (2.8 s mean duration, Extended Data Fig. 1e), while inactive bouts were considerably longer on the running wheel (7.3 s mean duration compared to 2.7 s mean duration in the VBA, Extended Data Fig. 1f). Consistent with our observations in the VBA, whisking decreased during the inactive state and increased slightly during the active state (Fig. 1k). These changes were accompanied by similar but not significant trend in pupil dynamics (Fig. 1l).

In summary, in both head-fixed contexts, mice spent most time in quiet wakefulness characterized by alternating active and inactive bouts, and less frequently initiated stereotyped behaviors such as egress, grooming, or running in a spontaneous, uniformly distributed fashion across session time. For subsequent analyses, we focused on egress, grooming, and running for two reasons: 1) we suggest that these stereotyped behaviors diverge from pure arousal fluctuations (unlike active and inactive states) and 2) these behaviors were rarer and long enough for a quiet baseline to be established before the transition, and therefore allow investigation of salient initiation of behavior. To this end, we selected only behavioral bouts that were preceded by a quiet baseline of at least 15 seconds. Specifically, for egress, we only included bouts in which no other egress event longer than 1 second occurred within the preceding 15 seconds. Similarly, for running, we selected bouts with no other running episode longer than 5 seconds within the same pre-bout interval. This ensured that we were capturing transitions from relative quiescence into the behavior of choice.

### Behavioral transitions can be predicted from motion, whisking, and pupil size shortly before onset

We next asked whether and when transitions into stereotyped behaviors can be predicted from behavioral readouts alone. To test this, we trained linear SVM classifiers (50 SVMs, 10-fold cross-validation) for each time point around behavior onset to distinguish transitions into egress, grooming, and running from pseudo-bouts (Fig. 1m). Pseudo-bout data was generated by extracting time windows (12 seconds before and after) around real transitions from shuffled sessions. This approach allowed us to create a control condition with the same bout statistics but without aligned behavioral onsets.

To assess when upcoming transitions became decodable, we computed the earliest decoding time (EDT) as the first time point (moving from behavior onset backward in time) at which decoding accuracy fell within one standard deviation above the mean control decoding accuracy^28^. This approach allowed us to identify the earliest moment at which behavioral states could be predicted from motion, whisking, and pupil data (Fig. 1n).

For egress, decoding from burrow position, whisking, and pupil size revealed an EDT of-0.5 to-1 second (Fig. 1n). Similarly, grooming was decodable from whisking 1.5 seconds before onset. Interestingly, pupil size predicted grooming above chance level up to 6 seconds in advance, and VBA burrow position 8.5 seconds (Fig. 1n). These longer EDTs likely reflect gradual behavioral shifts preceding grooming, such as an increase in pupil diameter and a change in the position in the burrow (Extended Data Fig. 2b). Aligning with these observations, when examining the behavioral state one frame prior to transitions into egress, grooming, and running, we found that active behavior increased above baseline immediately before the switch (Extended Data Fig. 3a-b).

Running could be decoded slightly above chance from slight movements of the wheel with an EDT of-21 seconds, whereas the EDT was-2 seconds from whisking and-1.5 seconds from the pupil (Fig. 1n). When comparing the average traces of behavioral variables for behavior bouts and pseudo-bouts, we found that wheel motion, whisking, and pupil size is at a lower baseline level than that of pseudo-bouts several seconds before the onset of running (Extended Data Fig. 2c). These results might suggest that on longer timescales mice are in a less active state prior to initiating running. By contrast, and consistent with previous reports for locomotion^35^, transitions from quiescence to movement on shorter timescales (0.5-2 s) are preceded by an increase in arousal, as reflected by enlarged pupil size and increased whisking (Fig. 1n, Extended Data Fig. 2).

Taken together, these results indicate that the initiation of egress, grooming, and running can be decoded from behavioral variables only a few seconds before onset, likely due to a short increase in arousal preceding the transition. Before that period, the transition cannot be decoded above chance, except for running, which is associated with slightly reduced locomotion.

### Distinct whole-brain activity patterns underlie egress, grooming, and running

While tracking behaviors in the VBA and running wheel, we simultaneously performed fUS to identify brain activity patterns associated with specific behaviors and their transitions. Although we focus on brain activity preceding behavior transitions, we first characterized the neural patterns underlying the stereotyped behaviors of interest.

Averaging voxel-wise fUS activity across multiple bouts revealed distinct whole-brain activity patterns for each behavior (Fig. 2a-c, Extended Data Fig. 4). Neural activity associated with running exhibited asymmetric activation, which is unlikely to reflect a biological effect, but rather the limited cranial window coverage across the cohort and the relatively low number of animals used (Extended Data Fig. 4c). Nevertheless, the brain-wide voxel data allowed us to decode behavioral bouts from pseudo-bouts in which the respective behavior was not exhibited, with high decoding accuracy using a similar SVM decoding approach utilized for the previous behavior decoding^31^ (Extended Data Fig. 4b).

**Fig. 2.**
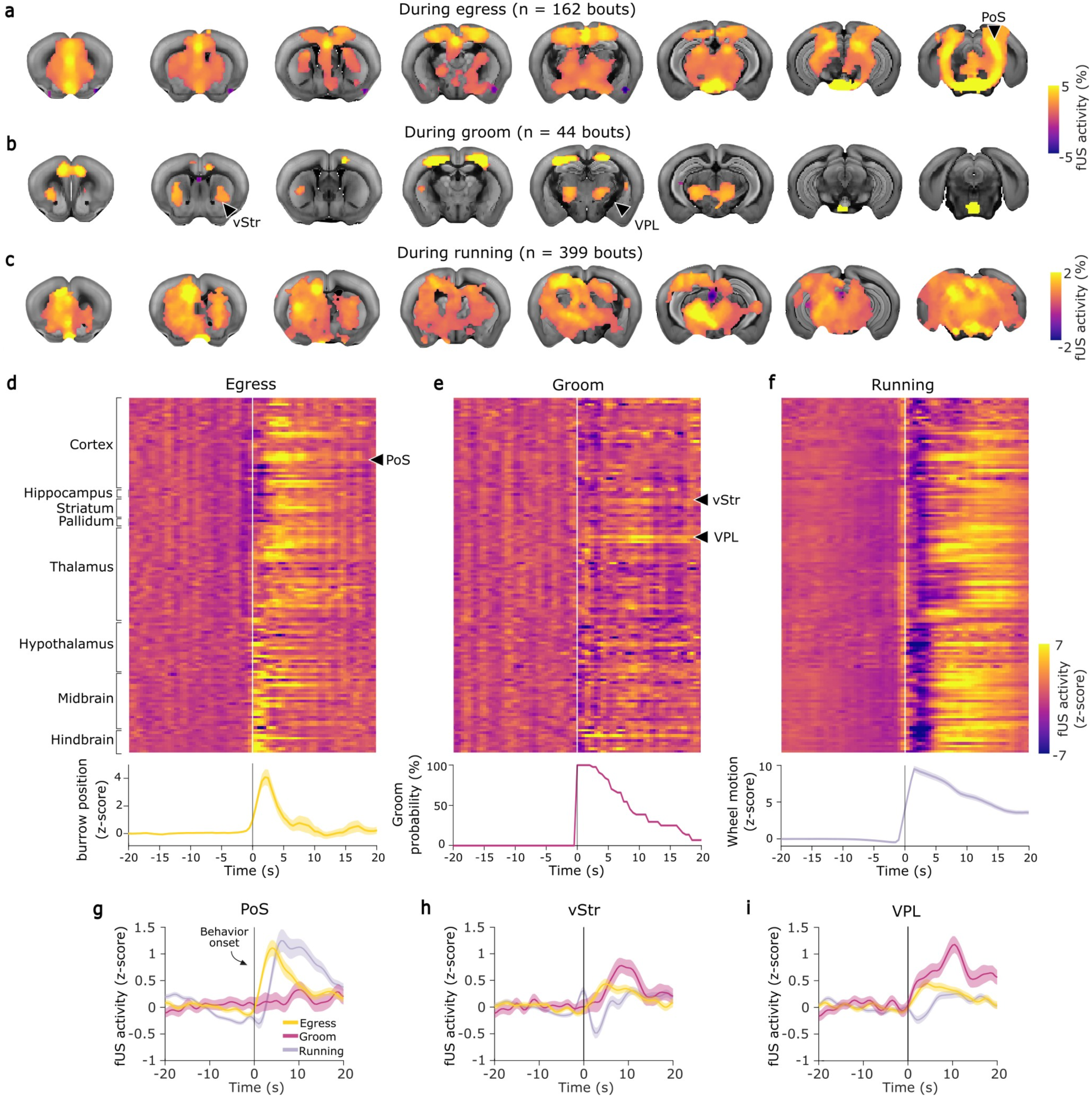
**Whole-brain activity patterns of egress, grooming and running. a-c**, fUS activity patterns underlying egress (**a**), grooming (**b**) and running (**c**) displayed on reference coronal brain images. Only voxels significantly different from zero are shown, FDR-corrected (FDR = 0.1), one-sample t-test, for egress adjusted p < 0.010, for grooming adjusted p < 0.0031, for running adjusted p < 0.017. Black arrows highlight regions shown in **g-i**: PoS = postsubiculum, vStr = ventral striatum, VPL = ventral posterolateral nucleus, **d-f**, top: per region fUS activity before and after onset of egress (**d**), grooming (**e**) and running (**f**). White line represents behavior onset for all panels. Bottom: corresponding behavior readouts. From left to right: burrow position, grooming probability and wheel motion. **g-i,** fUS activity of selected regions in PoS (**g**), vStr (**h**), and VPL (**i**) before and after onset of egress (yellow, mean ± s.e.m.), grooming (pink, mean ± s.e.m.) and running (gray, mean ± s.e.m.).

Across behaviors, we observed differences in region-wise neural dynamics after aligning and segmenting whole-brain data (Fig. 2d-f). Egress was associated with a wave of activity across many brain regions throughout the cortex, midbrain, thalamus, and striatum (Fig. 2d). As described before^37–39^, running also elicited activity in a broad set of regions throughout the midbrain, thalamus, and striatum, and widespread cortical activation at around 10 seconds after onset (Fig. 2f). Brain activity underlying grooming exhibited a sparser pattern of activity that was confined to a more selective set of regions, including the ventral striatum (vStr), ventral posterolateral thalamus (VPL), primary somatosensory cortex (SSp), pons, and prefrontal cortex, (Fig. 2b,e), consistent with previous studies^40,41^. In terms of temporal dynamics, the duration of the increased or decreased hemodynamic activity was consistent with the duration of the behavior itself, with egress eliciting shorter deviations from baseline (∼10 s) compared to running (∼20 s) and grooming (∼15 s) (Fig. 2d-f).

To visualize differences in activity across brain regions, we extracted signals of specific regions that showed significant activation in the whole-brain activity maps (Fig. 2a-c), and plotted their activity across time (Fig. 2g-i). Egress was associated with a rapid increase in activity in the postsubiculum (PoS) right after onset, whereas PoS activity remained unchanged during grooming and was elevated for a prolonged period after the onset of running (Fig. 2g). In contrast, the vStr and VPL showed a longer rise time after grooming onset, peaking at ∼10 seconds, and only small changes at the onset of egress and running (Fig. 2 h-i). These results illustrate the unique ability of fUS to reveal whole-brain activity correlates of stereotyped behaviors.

### Behavior transitions can be decoded from whole-brain activity several seconds in advance

Next, we examined whether changes in brain activity patterns precede transitions in behavior. To this end, we trained an SVM framework^42^ to differentiate fUS data recorded around the time of true behavior onsets from that of pseudo-bout onsets (Fig. 3a). As in the behavioral decoding analysis, we used only bouts with a quiet baseline period preceding the behavior to focus on transitions from quiescence to movement. Pseudo-bouts were selected using the same criteria, ensuring that mice were in the same behavioral state prior to onset in both conditions, thereby minimizing decoding bias (Fig. 3b).

**Fig. 3.**
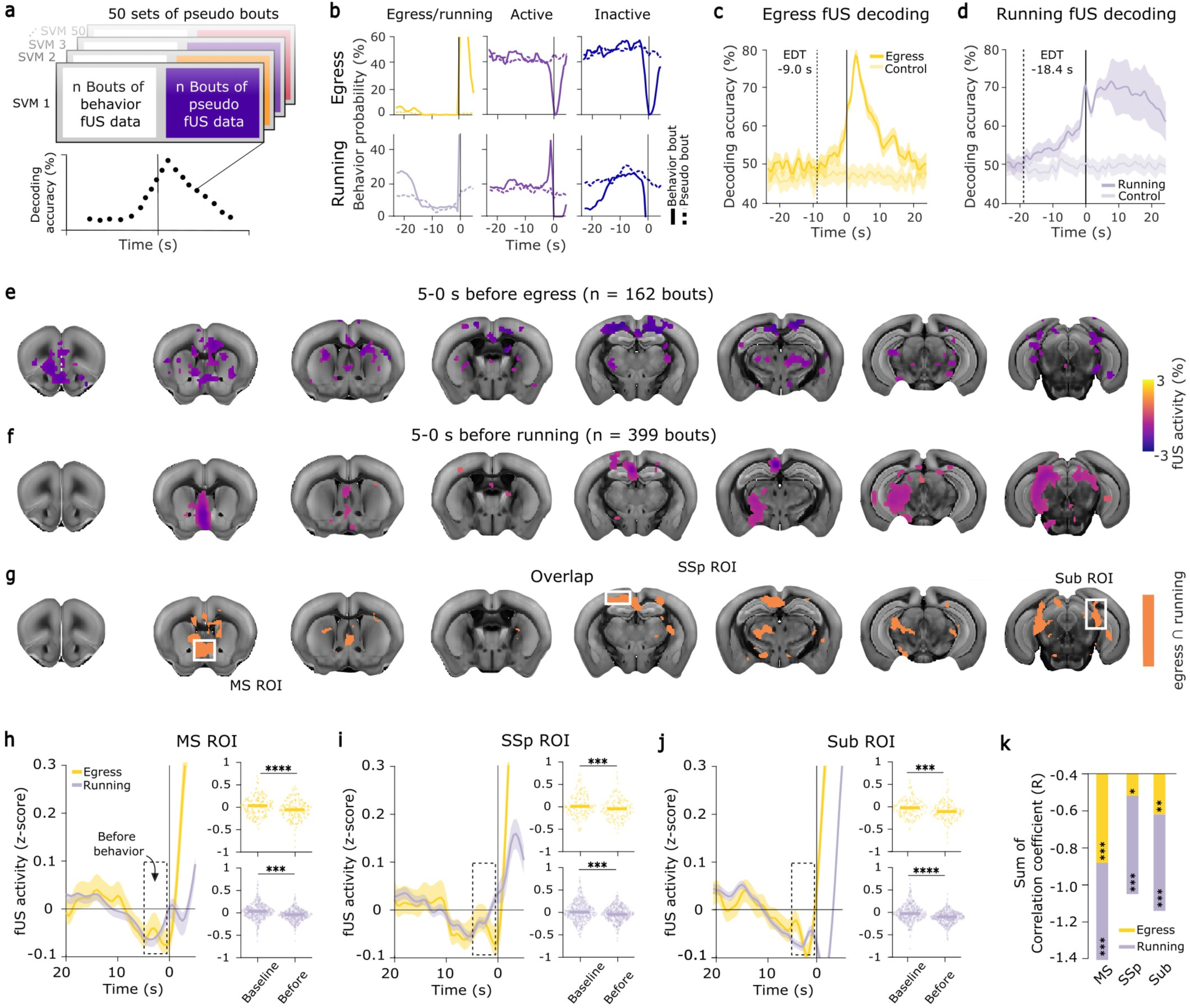
Behavior transitions to egress and running can be predicted from whole-brain data several seconds before. **a**, Schematic of whole-brain decoding using classification SVMs. Each data point represents the average decoding accuracy across 50 SVMs, run on fUS bout data and 50 different sets of pseudo-bouts (per run 10-fold cross-validation). **b**, Behavior state probabilities of behavior bouts (solid lines) around onset of egress (top) and running (bottom) and pseudo-bouts (dashed-lines). **c-d**, Whole-brain decoding (SVM classification behavior bouts vs. pseudo-bouts) using fUS data around egress (**c**, n = 162 from 9 animals here and elsewhere in the figure) and running (**d**, n = 500 from 3 animals here and elsewhere in the figure). Each timepoint represents an average decoding accuracy ± std. across 50 SVM runs each using a different set of pseudo-bouts. Control decoding accuracies (lighter yellow/gray) were generated by classifying one set of pseudo-bouts against 49 other sets of pseudo-bouts. Dashed lines indicate the EDT, defined as the first time point (moving from during behavior to before) at which the mean decoding accuracy (behavior bout vs. pseudo-bout) falls within one standard deviation of the control decoding accuracy (pseudo-bout vs. pseudo-bout). The EDT for egress (**c**) is-9.0 seconds and for running (**d**)-18.4 seconds. **e**, fUS activity patterns 5-0 seconds before the onset of egress on reference coronal brain images. Only voxels significantly different from zero are shown, uncorrected, p < 0.01, one-sample t-test. **f**, fUS activity patterns 5-0 seconds before the onset of running on reference coronal brain images. Only voxels significantly different from zero are shown, FDR-corrected (0.1), p < 0.004, one-sample t-test. **g**, Common significant regions before the onset of egress and running. Overlap of 20% most significant voxels depicted in orange. White boxes highlight common nodes (MS = medial septum, SSp = primary somatosensory cortex, Sub = subiculum). **h-j,** fUS activity of selected ROIs in MS (**h**), SSp (**i**), and Sub (**j**), before onset of egress (yellow line, mean ± s.e.m) and running (gray line, mean ± s.e.m). Dashed box highlights 5-0 seconds before behavior onset. Right: fUS activity during baseline episode and before egress (top, yellow) and running (bottom, gray). Horizontal lines show median fUS activity. Wilcoxon signed-rank test. **k**, sum of correlation coefficients (spearman’s rank correlation) of whole-brain decoding accuracies 10-2 seconds before behavior onset and fUS activity in ROI of MS, SSp, and Sub of egress dataset (yellow) and running dataset (gray) per session. * p < 0.05, ** p < 0.01, *** p < 0.001, **** p < 0.0001.

We found that transitions to egress were predictable from whole-brain activity 9 seconds before onset, with decoding accuracy gradually increasing before and peaking during the behavior (Fig. 3c). For running, decoding accuracy was significantly above chance as early as 18.4 seconds before onset, suggesting that distinct activity patterns emerge well in advance of the behavior transition. A more stringent EDT criteria did not change the fact that both behaviors could be decoded seconds before initiation from the brain data (Extended Data Fig. 6c-d). This long EDT is consistent with the decoding from wheel motion (Fig. 1n), but the gradual increase of decoding accuracy is only observed in the brain data and can therefore not be explained by behavioral changes alone. This indicates that there is a detectable change in brain state preceding a transition in behavior. Since the number of grooming bouts was limited (n = 44) and likely limited the statistical power of our decoding approach, we did not detect any significant (p < 0.05) brain activity before behavior onset and were not able to decode grooming before onset (Extended Data Fig. 5).

Next, we asked whether both the predictability and the associated whole-brain changes generalized across behaviors. We hypothesized that a common brain state may precede and potentially facilitate spontaneous transitions from quiescence to different behaviors. To this end, we first averaged voxel-wise fUS activity 5 seconds before the onset of egress and running and found that several brain regions exhibited a decrease in activity prior to both behaviors (Fig. 3e-g, Extended Data Fig. 6). To uncover regions that might be part of a general network associated with behavioral switching, we superimposed significantly modulated voxels for both egress and running. We identified several key regions including the MS, SSp, and subiculum (Sub) (Fig. 3g) that are down modulated long before the transition, and that may reflect a transition-prone brain state. When looking at a shorter time window before behavior initiation, we found that periaqueductal gray (PAG) activity increased around 2 seconds before both behaviors, suggesting a more direct role in initiating the transition itself (Extended Data Fig. 7).

When examining the activity of regions significantly modulated before behavior onset for both egress and running, we found that they (MS, SSp, and Sub) showed a gradual decrease in activity starting around 10 seconds before onset, and which continued until approximately 2 seconds before behavior initiation (Fig. 3h-j). When correlating the whole-brain decoding accuracy with the signals of identified key regions, we found that decreasing signals in MS, SSp, Sub, and PAG co-occur with increasing predictability in the 10 to 2 seconds preceding behavior initiation for both egress and running (Fig. 3k, Extended Data Fig. 8). This finding might suggest that a reduction in brain activity in certain areas permits or facilitates behavioral transitions or, more generally, that a brain state that facilitates behavioral transitions involves the inhibition of these regions. Notably, the strongest correlation among identified candidate regions was observed in the MS (Fig. 3k). This pattern shifted in the final two seconds before onset where increasing fUS activity in these regions coincided with increasing decoding accuracy (Extended Data Fig. 9). This timing is associated with increases in whisking, motion, and pupil size (Fig. 1), pointing to a rising arousal state that may accompany behavioral execution. Taken together, these findings highlight a dynamic two-stage process: 1) an early reduction of activity in specific regions, followed by 2) a short arousal-driven increase of activity that may govern behavioral initiation.

### Optogenetic inhibition of MS neurons increases the likelihood of behavioral transitions

To investigate whether the decrease in activity observed before behavior initiation is merely correlated with behavioral transitions or causally implicated in switching itself, we inhibited the MS using optogenetics. Specifically, inhibition was performed for a prolonged period to mimic the observed decrease in brain activity and to assess its behavioral impact (Fig. 4a). We selected the MS as our target because it emerged as a prominent node in the superposition map of significant voxels preceding both egress and running (Fig. 3g), and its decrease in activity was associated with increasing decoding accuracy in the pre-transition window (Fig. 3k). Moreover, the MS has been previously implicated in arousal and behavioral state transitions^12,43,44^, aligning with our hypothesis that its activity could play a causal role in mediating behavioral transitions.

**Fig. 4.**
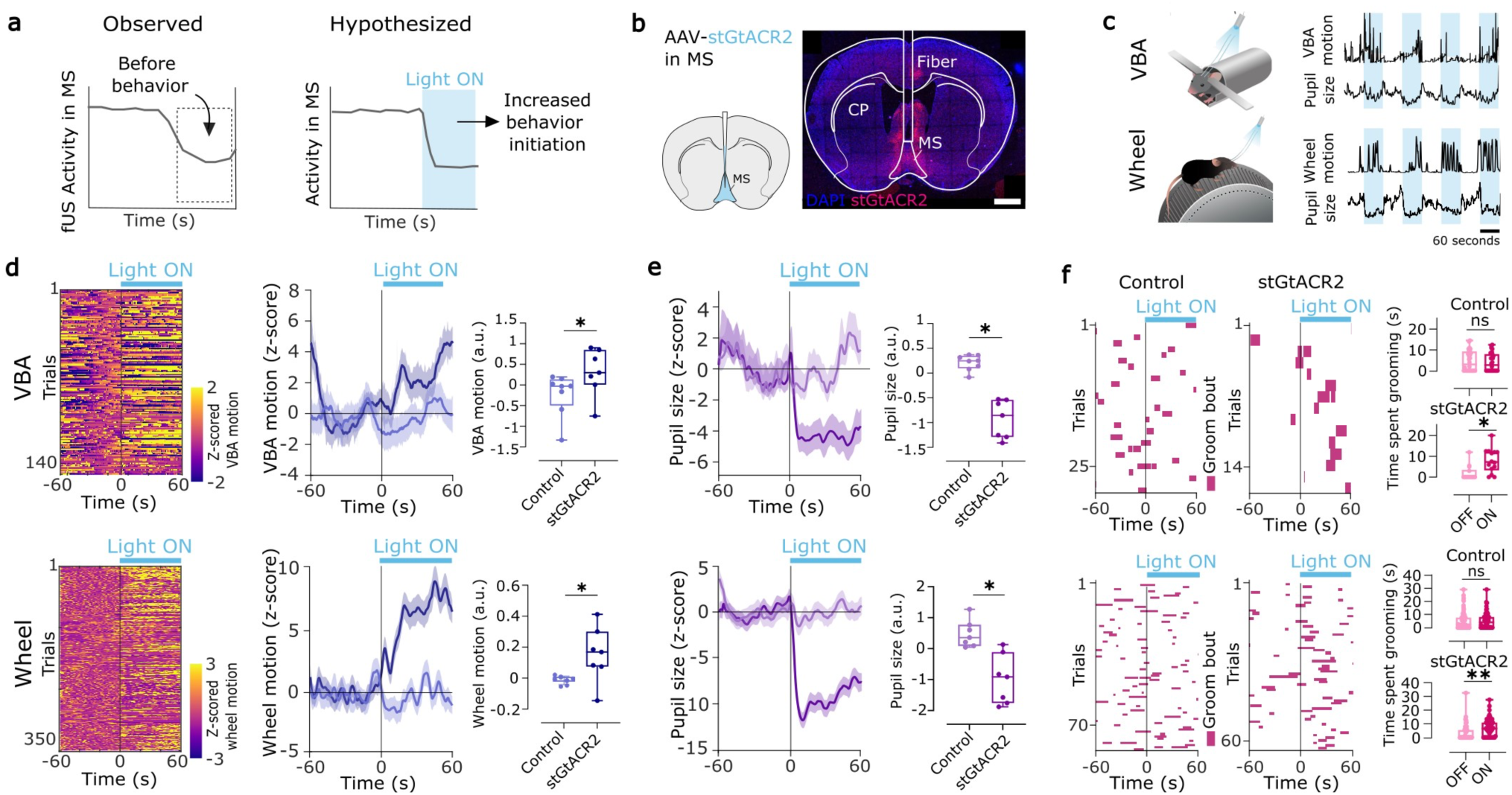
Inhibition of medial septal neurons causes an increase in movement in the VBA and on the running wheel and enhanced grooming. a, Schematic of hypothesis underlying optogenetic experiment. Left: Observed fUS decrease in MS before behavior onset. Right: hypothesis that decreasing MS activity via inhibitory optogenetics might increase behavior initiation. b, Injection of pAAV-CKIIa-stGtACR2-FusionRed and optic fiber implantation. White scale bar represents 1 mm. c, Representative examples of two different mice showing the effect of light stimulation on motion and pupil size in the VBA on top, and on wheel motion and pupil size below on the running wheel. d, VBA Motion (top) and wheel motion (bottom) before and during light stimulation (for VBA comparing first 30 seconds of light stimulation of control (n = 8 mice, pAAV-CaMKIIa-mCherry) to stGtACR2 (n = 7 mice), for wheel comparing motion during whole 60 second stimulation period of control and stGtACR2, dark blue stGtACR2, light blue control, mean ± s.e.m., Wilcoxon rank sum test, p < 0.05). e, Pupil diameter before and during light stimulation in VBA (top) and on running wheel (bottom) (comparing pupil size of control and stGtACR2 for whole 60 second stimulation period, dark purple stGtACR2, light purple control, mean ± s.e.m., Wilcoxon rank sum test, VBA p< 0.001, Wheel p < 0.01. f, Groom occurrence before and during inhibition of MS neurons in VBA (top) and on running wheel (bottom). Quantified length in seconds per trial and compared on and off periods for control and stGtACR2 groups (Wilcoxon matched-pairs signed-rank test, VBA: n = 29 trials from 8 control mice (p = 0.731) and n = 15 trials from 7 stGtACR2 mice (p = 0.019) in the VBA, n = 75 from 8 control mice (p = 0.816) and n = 63 from 7 stGtACR2 mice (p = 0.0011) on the wheel for stGtACR2. * p < 0.05, ** p < 0.01, *** p < 0.001.

For this purpose, we expressed the soma-targeted anion-conducting channelrhodopsin (stGtACR2) in CaMKIIa-positive neurons in the MS of 7 mice (Fig. 4b). Transcriptomic data shows that many cell types in the MS, including cholinergic cells in the MS express CaMKIIa^45^. As a control, we expressed mCherry in CaMKIIa-positive cells in the MS of 8 mice in an independent cohort. Next, we activated the opsin for 60 seconds using continuous blue (473 nm) light stimulation in the VBA and on the running wheel and subsequently quantified its behavioral outcome (Fig. 4c).

Prolonged inhibition of CaMKIIa-positive neurons in the MS led to an increase in movement in the VBA during the first 30 seconds of light stimulation and on the running wheel throughout the entire 60-second stimulation period (Fig. 4d). Additionally, we observed a pronounced decrease in pupil size in both contexts during MS inhibition (Fig. 4e). To further assess how this manipulation influenced behaviors, we calculated the probabilities of egress, grooming, and running behaviors before and during light stimulation. Comparing the change in behavior probabilities before and during inhibition to that of a shuffled distribution revealed that the likelihood of all three behaviors increased significantly during MS inhibition (Extended Data Fig. 10a-d).

We further analyzed grooming behavior in the opsin and control cohorts and found that mice expressing stGtACR2 spent significantly more time grooming during inhibition in both experimental contexts, while no such change was observed in control mice (Fig. 4f, Extended Data Fig. 10e). Importantly, this increase was not due to longer bouts, but rather to a greater number of groom bouts during inhibition (Extended Data Fig. 10f).

These results show that inhibiting the CamKIIa-positive neurons in MS increases the probability of mice initiating different behaviors. Taken together, these findings suggests that reduced MS activity observed via fUS is not merely a correlate of behavioral switching, but may causally contribute to facilitating transitions. By reducing activity in the MS, the brain may enter a transition-prone state that increases the likelihood of switching behaviors.

## Discussion

This work reveals a brain-wide hemodynamic pattern that predicts uninstructed behavior transitions in mice. A coordinated decrease starting several seconds before behavior onset in MS, Sub, and SSp makes transitions predictable, and might prime the brain for state switching. Optogenetic inhibition of the MS, mimicking this observed decrease, increased the likelihood of initiating behaviors such as locomotion and grooming, identifying the MS as a potential key hub in the facilitation of spontaneous behavioral transitions.

Brain signals preceding voluntary actions have been observed in cortex via EEG as early as in the 1960s^3^. Notably, it was observed that there is a point in time around 200 ms before the execution of movements, when it cannot be stopped by the participant^46^. While this finding demonstrates that a causal process begins hundreds of milliseconds before action, it also indicates that the preceding signal is not strictly deterministic. In our work, noticeable predictability began roughly 10 seconds before the initiation of egress and running, yet in line with this finding, we interpret this not as evidence that the brain has fully committed to a specific action, but rather that it has entered a state in which initiating action becomes increasingly probable.

More recently, researchers have employed machine learning to determine the time point at which a decision can be decoded from fMRI data. It was found that upcoming choices to press a left or right button could be predicted from activity in the parietal and frontal cortex up to ∼7 seconds before the action^5^. This study demonstrated that predictive information about decisions is present in the brain several seconds prior to movement initiation, a finding that aligns with our own results. A more recent study adopted a similar approach in mice, recording calcium signals as a proxy for neural activity across the dorsal cortical surface while animals performed self-initiated lever pulls^28^. It was reported that uninstructed movements can be predicted 3-5 seconds before onset. Our observed EDTs of 18 seconds for running and 9 seconds for egress might be explained by the fact that we have data from the whole mouse brain, which might make earlier decoding possible, and reveals that the brain might enter a transition-prone state much earlier.

It has been suggested that this gradual increase in decoding accuracy prior to uninstructed movement initiation may be explained by leaky stochastic accumulator models^47^. These accumulator models, traditionally applied to perceptual decision-making, describe how sensory evidence is integrated over time until a decision threshold is reached, triggering an action^48^. Recently, several studies have speculated that leaky accumulator models may also account for spontaneous behavior transitions ^28,47–49^. Our results align with these proposals, and we suggest that the steady increase in decoding accuracy several seconds before transitions reflects the accumulation of noisy fluctuations and internal drive that eventually push the system across a decision boundary, resulting in a behavioral switch.

Having access to almost the entire mouse brain via fUS, we found an overlap of voxels significantly modulated before egress and running. We identified the MS, SSp, and Sub as the main hubs of this overlap. We speculate that this decrease across regions might reflect a shift in internal state that makes the animal more likely to initiate transitions. In humans, the SSp specifically has been shown to predict upcoming movements several seconds before onset, suggesting that this region changes its processing state well in advance, and potentially prepares for upcoming sensory information^50^. This finding could similarly reflect preparation for new somatosensory inputs when transitioning between behaviors. Similarly, the Sub, which has been implicated in spatial navigation^51–53^, might enter a distinct processing state in preparation for new location and motion information when behavior changes. Moreover, a recent study reported that excitation of several projections to the ventral Sub promotes wakefulness and arousal in mice^54^. Accordingly, a reduced baseline activity in the Sub may reflect a low-arousal yet transition-prone state, increasing the likelihood of major behavioral changes. Consistent with this idea, we find that mice are more likely to initiate locomotion from an extended period of lower arousal.

fUS measures changes in cerebral blood volume, from which neural activity can only be inferred indirectly. However, several studies have demonstrated that fUS intensity correlates strongly with direct measures such as electrophysiology^55–57^, providing confidence that the observed hemodynamic patterns reflect bulk neural activity. This correlation is valid for both increases and decreases in blood volume, such that decreases in fUS signals were shown to reflect inhibition of the neural population^57^. Finally, head-fixed experiments limit the breadth of behaviors that mice can perform, but volumetric fUS in this context still provides the best trade-off between brain coverage and behavioral complexity ^58^. Moreover, head fixation is necessary for volumetric fUS due to the bulkiness of the matrix ultrasound probe used to capture such a large field-of-view. Future work could extend this framework to other behaviors to assess whether the processes identified here generalize, potentially including freely moving paradigms using a single-plane fUS imaging, at the cost of much less brain coverage.

To determine whether the reduction in MS activity identified by fUS reflected a causal mechanism rather than a mere correlation with behavioral transitions, we used optogenetics to broadly inhibit cells in this region. We found that mimicking the naturally observed decrease in fUS activity, increased the likelihood of initiating spontaneous behaviors such as locomotion and grooming. These findings suggest that a prolonged reduction in MS activity facilitates behavioral transitions, potentially by enhancing the drive to switch behaviors. Based on these results, we speculate that the MS mediates a general increase in behavioral flexibility rather than simply an increased tendency towards movement. This is specifically supported by the fact that we see not only an increased incidence of running on the wheel and of motion in the VBA, but also more grooming bouts in both setups during optogenetic inhibition of MS.

The MS is both a cholinergic hub within the basal forebrain and a major input to hippocampus, where it is best known for regulating hippocampal theta oscillations^59–61^ that serve distinct behavioral purposes. These oscillations have been described to increase shortly before the onset of locomotion and are thought to bring the hippocampus into a processing-ready state for upcoming spatial information^59,60^. Specifically, a population of glutamatergic MS neurons has been found to entrain theta oscillations in the hippocampus and control the initiation of running and its velocity^61^. Moreover, it was recently found that a glutamatergic projection from the MS to the ventral tegmental area (VTA) drives exploratory behaviors^62^. In our study, inhibition of MS neurons led to an increase in behavior initiations, which suggests that we did not target primarily glutamatergic neurons. Transcriptomics data of the MS shows that CaMKIIa-positive neurons in the MS, among others, consist of cholinergic neurons^45^, which might lead to the activation of the glutamatergic population within the highly interconnected MS when inhibited^63,64^. Moreover, a recent study reported increased locomotion and exploration following chemogenetic inhibition of cholinergic neurons in the MS, consistent with our observations. We also found that inhibition of MS neurons led to strong pupil constriction, in line with previous reports linking pupil dilation to increased cholinergic activity^65^. Follow-up work will be needed to decipher the effect of inhibition on different cell types and projections.

According to leaky accumulator models^47,48^ and the hydraulic model^66,67^, the probability of transitioning behaviors increases with the duration of an action not being performed, and leads to a buildup of internal drive. Within this framework, we propose that the prolonged reduction in MS, SSp, and Sub activity, as well as the increase of decoding accuracy about 10 seconds before behavior onset, reflects such a pre-initiation state. During this state, internal drive accumulates and progressively biases the animal toward behavioral switching. This transition-prone state is followed by a brief arousal increase immediately preceding movement onset, marked by increases in whisking, pupil diameter, and body motion.

Overall, using whole-brain fUS during spontaneous behavior in mice, we find that distributed hemodynamic patterns predict behavioral transitions well before their execution. Our results indicate that seemingly spontaneous behaviors do not in fact emerge spontaneously, but are more likely to emerge from a transition-prone internal state that accumulates over seconds and which may be driven by a network of brain regions, with the MS acting as a key node.

## Methods

### Animals

All animal experiments and care were performed in accordance with the guidelines of the Max Planck Society, the government of Upper Bavaria, and Lower Saxony State Office for Consumer Protection and Food Safety (LAVES). Surgeries were performed on male and female mice aged between 8-10 weeks old and maintained in a C57Bl/6N background. The mice were group housed in ventilated cages in a 12-hour light/dark cycle and fed ad libitum.

### Cranial window implantation

As the skull greatly attenuates ultrasound waves, we performed fUS through a large chronic cranial window. To this end, mice were first anesthetized with a subcutaneous injection of an anesthetic cocktail consisting of fentanyl (0.05 mg/kg), medetomidine (0.5 mg/kg) and midazolam (5.0 mg/kg) (FMM). The depth of the general anesthesia was confirmed by the absence of typical reflexes (righting, eyelid and interdigital reflexes). Mice were then secured using a bite bar and placed on a heating pad (Supertech) at 37°C. The animals’ eyes were protected from drying out with Bepanthen eye ointment (Bayer), the hair was removed over the dorsal skull, and the skin was disinfected. A midline skin incision was then made over the skull to expose the bone, which was cleaned with saline. A dental drill with a 0.5 mm burr (Hager & Meisinger) was then used to make a large cranial window into the skull, after which most of the dorsal skull was removed using forceps while keeping the dura intact. A custom cranial implant, the COMBO window^68^ was then attached to the peripheral skull using cyanoacrylate glue (Pattex) and dental cement (Super-Bond). Before antagonizing the anesthesia with a subcutaneous injection of flumazenil (0.5 mg/kg) and atipamezole (2.5 mg/kg), buprenorphine (0.1 mg/kg) was injected for pain relief. For post-surgery analgesia, a subcutaneous injection of buprenorphine (0.1 mg/kg) and carprofen (5 mg/kg) is provided once a day for at least 48 h, and until no more pain symptoms can be observed. Several days after the cranial window implantation, mice were again anesthetized using FMM to implant a metal head plate for later head fixation in the VBA and on the running wheel. After the recovery period, mice were progressively habituated to the experimenter and to the recording setup.

### COMBO window

The COMBO window is a 3D-printed polymer implant designed to enable chronic imaging through a large cranial window in mice. It was developed to provide long-term stability, access to a substantial portion of the brain, and compatibility with multiple modalities, including functional ultrasound, acute electrophysiology, calcium imaging, and optogenetics^68^. After 3D-printing the black polymer implant using a Formlabs printer, a protective polymer film (GoodFellow) was glued to the inner surface of the implant using Superglue (Pattex) and Epoxy (Thorlabs). This film later covers the exposed brain and shields it from external influences. Following implantation, a second short, non-invasive surgery was performed to securely attach the head plate directly to the COMBO implant. For this study, we used a bilateral version of the COMBO window, allowing for stable bilateral head-fixation in both the VBA and running wheel setups.

### Virtual burrow assay

The VBA is a setup in which mice are placed and head-fixed in an air-floating tube which is movable. By pushing the burrow back, mice can voluntarily exit the burrow in a movement called egress. The VBA was designed to mimic the natural burrowing behavior of mice in head-fixed conditions, allowing behavioral and neuronal recordings with little habituation and training^33^. The VBA consists of a 3D-printed burrow that is air-lifted along metal rails to allow for smooth movement. The burrow is connected via a tether to a force sensor, which measures the force exerted by the mouse when retreating into the burrow. In addition, a laser displacement sensor records the movement of the burrow which is used for detecting egress. A small object (e.g., metal screw, plastic cap) was attached to a connector on the burrow to encourage more egress events. Importantly, this object does not change throughout the session. Head-fixation was achieved using bilateral head bars connected to head plates that were secured to the COMBO window implants. The setup also included two FLIR Blackfly S cameras (Fig. 2.1a) recording at 30 Hz: one positioned to capture full-body movement and the other focused on the face to enable quantification of whisking and pupil size. A custom probe holder (Thorlabs) secured the 3D fUS probe above the cranial window for whole-brain imaging. Illumination was provided by three infrared LED arrays (Kemo Electronics). A small adjustable reading-lamp directed toward the face of the mouse was used to maintain constant mild illumination inside the behavior box. The cameras, fUS recording, and VBA were all synchronized and triggered using a TTL-pulse generator (OPTG-8, Doric lenses). Before data acquisition, mice were handled for three consecutive days (at least 10 minutes per day) to habituate them to the experimenter. This was followed by a short 5-minute habituation session inside the VBA. Subsequent recording sessions lasted 10 minutes per day (5-10 sessions per mouse), during which the environmental conditions inside the behavior box remained constant.

### Running wheel

To record whole-brain activity via fUS during spontaneous locomotion, we used the KineMouse Wheel setup^69^. The system includes a centrally mounted running wheel and a custom 3D probe holder (Thorlabs) that allows stable positioning of the fUS probe on top of the mouse brain during running. Two FLIR Blackfly S cameras recording at 20 Hz were integrated into the setup, one to record body movements and the other to capture facial features for subsequent analysis of whisking and pupil size via Facemap^34^. Illumination was provided by a constant ambient light source and two infrared LED arrays. The cameras and fUS recording were simultaneously triggered using a TTL-pulse generator (OPTG-8, Doric lenses). Prior to recordings on the running wheel, mice were handled by the experimenter for 3 consecutive days, 10 minutes per day. This was followed by a 5-day wheel training protocol, during which mice were gradually habituated to the setup. Habituation sessions increased in duration across days: 5 minutes on day 1, 15 minutes on day 2, 30 minutes on day 3, 45 minutes on day 4, and 60 minutes on day 5.

### fUS acquisition

Before starting fUS acquisition, the ultrasound probe was mounted onto the probe holder in either the VBA or running wheel setup and positioned to align with the center of the cranial window. The probe position was verified by acquiring Doppler images and/or B-mode images in real-time, which were used to guide fine adjustments in probe positioning to ensure optimal coverage and alignment. fUS recordings were performed using a 32 × 32 channel volumetric probe (15 MHz, Vermon) with a spatial resolution of 220 × 280 × 175 μm^70^ connected to a Vantage 256 (Verasonics, Inc.) using a 4x multiplexer connector (UTA 1024 MUX, Verasonics, Inc) and controlled by a custom fUS acquisition module for 3D fUS imaging (AUTC). The beamforming and acquisition sequences were adjusted accordingly. To generate one doppler image, 160 compound images were captured at a pulse repetition frequency of 400 Hz and incoherently averaged. A compound ultrasound image was generated by summing plane wave emissions at-4.5,-3,-1.5, 0, 1.5, 3, and 4.5 degrees. Real-time clutter filtering was applied using SVD and the first 20% of singular vectors were discarded. This processing pipeline yielded one power Doppler image approximately every 500 milliseconds. Due to improvement of the computational power over time, the acquisition time was different for the VBA cohort (0.5 s) and for the wheel cohort (0.4 s).

### Stereotaxic surgery

Viral injections and implantations of optic fibers were performed using a manual stereotaxic apparatus (Stoelting). Mice were anaesthetized, using FMM in the same concentration as for cranial window implantation, secured in the stereotaxic frame, and their temperature was maintained using a heating pad. For optogenetic inhibition mice were either injected 500 nl of pAAV-CKIIa-stGtACR2-FusionRed (Addgene, #105669-AAV1, 1×10^13 vg/mL)^71^ or pAAV-CaMKIIa-mCherry (Addgene, #105669-AAV1, 1×10^13 vg/mL) as control in the MS (AP: 0.98, DV: 4.25, ML: 0) at a rate of 50 nl/min. To prevent backflow, the pipette was kept in place for 10 minutes after each injection. After virus injection, optic fibers (200 μm core, 0.22 NA, Thorlabs) were implanted 200 μm above the injection site using Superglue and dental cement. A COMBO window (without the polymer film, since no cranial window was made) was mounted onto the skull to allow attachment of the head plate and later head fixation in the VBA and running wheel setups. Before reversing anesthesia, mice received a subcutaneous injection of buprenorphine (0.1 mg/kg) for postoperative analgesia. Anesthesia was then antagonized with subcutaneous injections of flumazenil (0.5 mg/kg) and atipamezole (2.5 mg/kg). For post-surgery analgesia, a subcutaneous injection of buprenorphine (0.1 mg/kg) and carprofen (5 mg/kg) is provided once a day for at least 48 h, and until no more pain symptoms can be observed. Several days later, mice were re-anesthetized using FMM to surgically implant a metal head plate, allowing for stable head fixation during recordings in both behavioral setups. Experiments were performed starting after 3 weeks expression time.

### In vivo optogenetics

Implanted optical fibers in the MS were connected to a 473-nm laser (CNI) via metallic mating sleeves (Thorlabs) to activate the opsin stGtACR2. Before starting experiments, laser power was set to 5 mW using a photometer (Thorlabs). The stimulation was controlled using a programmable TTL pulse generator (OPTG-8, Doric lenses). Stimulation consisted of 60-second continuous light stimulation periods followed by 60-seconds no light stimulation periods. The order of stimulation was alternated every other day, starting with either an on or off period. Optogenetic sessions in the VBA lasted 10 minutes, while they lasted 30 minutes on the running wheel.

### Histology

To verify virus expression and confirm the correct placement of optical fibers, histological analysis was performed after the completion of behavioral experiments. Mice were anesthetized with a mixture of ketamine and xylazine and transcardially perfused with approximately 25 ml of 0.1 M phosphate-buffered saline (PBS, Life Technologies), followed by 25 ml of 4% paraformaldehyde (PFA, Thermo Scientific) in PBS. Afterward, brains were post-fixed in 4% PFA overnight at 4 °C. The following day, brains were rinsed in PBS and incubated in a 27.4% sucrose solution (Sigma-Aldrich) in PBS at 4 °C. Brains remained in the sucrose solution until they sank to the bottom of the tube, indicating full sucrose saturation for optimal cryo-protection. Next, the tissue was rinsed in PBS once, dried carefully with tissue and embedded in section medium (NEG-50, Epredia) in cryo-molds (Sakura) and stored at - 20 °C. A Cryostat (Leica) was used to acquire 25-μm-thick coronal brain slices, that were directly mounted onto adhesion slides (Superforst Plus, Thermo Scientific). Brain slices were stained with 20 mM Hoechst Solution (#62249, Thermo Scientific) or directly embedded using antifade mounting medium (#H1220, Vectashield). Brain sections were imaged using a Leica STELLARIS confocal microscope using a 20x air objective (Leica Microsystems). LAS X (Leica Microsystems) and ImageJ were used to process the acquired images.

### Behavior classification

All behavioral and fUS data were analyzed using MATLAB (MathWorks) or Prism 10 (GraphPad Software). The significance level, along with the number of mice, sessions, or trials, are reported in the respective figure legends. Detailed descriptions of the analysis procedures and the statistical tests used are provided below.

### Active / Inactive

To classify behavioral states as active or inactive, a readout of whisking was extracted from facial video recordings. A region of interest (ROI) was manually selected from the whisker pad of each mouse, and motion signals were extracted using SVD in Facemap^34^. Facemap computes the dominant motion component within the ROI, resulting in a single trace that reliably reflects whisking activity. This trace and all other behavior data was further processed using custom MATLAB scripts. For classification, a threshold was defined as a multiple of the standard deviation of the motion SVD trace (2× for motion SVD data from the VBA setup and 1.3× for motion SVD data from the wheel data). Values above the threshold were labeled as active, and those below it as inactive. A moving average with a 0.16-second window and mode filter with a 0.1-second window were applied to the resulting binary signal to smooth transitions and reduce noise.

### Egress

To identify egress events from the burrow in the VBA setup, we analyzed the voltage signal from the laser displacement sensor. The raw voltage signal was first converted into burrow displacement in millimeters (mm) using a linear transformation based on sensor calibration and servo position:

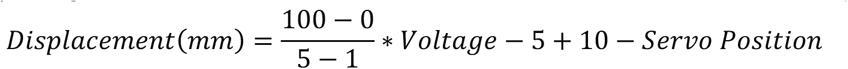

We then applied a sliding window of 10 samples (corresponding to 1 second of recording) to detect egress. If the displacement range within this window exceeded a threshold of 14 mm, the corresponding time point was classified as egress. This approach ensured that only sharp and strong exits from the burrow were marked as egress. For further analysis, we included only events that lasted longer than 1 second. To ensure accurate timing of bouts, egress and groom bouts were manually checked after automatic detection.

### Groom

For groom classification, a ROI in the snout area of the mouse was manually defined for each session. Next, the first 100 frames (corresponding to 5 seconds) of the ROI were averaged to create a baseline frame. This baseline was then correlated with every subsequent frame in the video to quantify changes in the ROI over time. Frames whose mean correlation with the baseline dropped below a threshold of 0.3 were classified as groom, based on the observation that grooming causes large changes in the snout ROI due to paw movements, leading to a substantial drop in correlation. All automatically detected groom bouts were manually curated: their onset timing was adjusted if needed, and false positives were removed from the dataset. In contrast to the VBA dataset, where 44 groom bouts were identified, neither automatic detection nor manual inspection revealed any groom bouts in the running wheel dataset.

### Running

To classify running in the wheel dataset, an ROI was manually selected on the running wheel from the body camera recordings. For each frame, the absolute difference from the subsequent frame within the ROI was computed to estimate motion energy. This metric reflects the movement of the wheel where high values indicate running and low values correspond to periods of inactivity. Time points where motion energy exceeded 2 times the standard deviation of the session, were classified as running. To reduce noise, the motion energy signal was smoothed using a moving average with a 0.35-second window, followed by mode filtering with a 0.15-second window. Consequently, the temporal precision for detecting the onset of behavioral bouts is 0.35 seconds. All behavioral labels were combined hierarchically, such that the more distinct behaviors of grooming, egress, and running overruled baseline classifications of active and inactive states. For fUS analysis, we included only running bouts that lasted longer than 5 seconds.

### Behavior quantification

To calculate the occurrence of each behavioral state, the total number of time points labeled as a given behavior across all sessions was summed, divided by the total number of time points, and multiplied by 100 to obtain the percentage of time spent in each state. These percentages were visualized as a pie chart using MATLAB. To visualize behavioral readouts in Fig. 1 aligned to the onset of different behavioral states - active, inactive, grooming, egress, and running - a trial-averaged trace was computed. Data were z-scored to a 30-second baseline for VBA recordings or a 24-second baseline for wheel recordings. The resulting traces were plotted as the mean ± standard error of the mean (SEM). For animal-wise quantification, behavioral readouts (z-scored to baseline) during each behavioral state were averaged across animals. To compare differences between behavioral states, Restricted Maximum Likelihood calculations were performed for mixed model analysis followed by Tukey’s post hoc multiple comparisons.

### Behavior decoding

To assess whether motion, whisking, or pupil size could predict behavioral transitions, we trained linear SVM classifiers with a custom written script in MATLAB to distinguish behavior bouts from pseudo-bouts. True behavior bouts were defined as 10 seconds of data preceding and 5 seconds following the onset of egress, grooming, or running transitions. By contrast, pseudo-bouts were generated by taking timestamps of true behavior onsets from shuffled session identities, generating a control bout set that differed only in the fact that the transitions were not aligned. For both bout sets the readouts were z-scored to the whole pre-onset (10-second) baseline. Note that our behavioral variables were downsampled to 2 Hz for the VBA data and 2.5 Hz for the wheel data to match the fUS sampling rates. We generated 50 independent pseudo-bout sets and trained 50 linear SVM classifiers per time point using 500 real and 500 pseudo-bouts per iteration. This allowed us to compute the mean decoding accuracy over 50 runs. Additionally, each classifier was evaluated using 10-fold cross-validation. As a control, we also computed decoding accuracy from 50 SVM runs comparing one pseudo-bout set to 49 different pseudo-bout sets. The resulting mean and standard error of decoding accuracy were plotted in Fig. 1. Inspired by Mitelut et al^28^, we next determined the earliest decoding time (EDT), which we defined as the first time point, preceding the behavior onset, at which decoding accuracy (behavior vs pseudo-bout) increased beyond one standard deviation above the mean control (pseudo vs pseudo-bout). This allowed us to determine the time at which upcoming behaviors became predictable above chance.

### fUS preprocessing

The fUS data was preprocessed using custom-written MATLAB scripts. First the data was temporally interpolated to correct for timing differences in fUS acquisition and to obtain a consistent acquisition time of 500 or 400 ms. Next, the brain volumes were registered to a reference mouse brain atlas (Allen Brain Institute). For each mouse, the registration was performed manually for the first imaging session via a GUI that allows alignment of an average power Doppler image to the atlas^31,32^, and subsequent sessions were registered to the first one automatically using a custom code. As differences in brain coverage may occur between mice due to cranial window coverage, bone/dura regrowth, etc., a brain mask was hand-drawn (ITK-SNAP software) to ensure that only high quality fUS data was included. The Allen Brain Insitute’s atlas was used to determine the limits of the brain. Both masks were combined to determine the final brain coverage mask. Next, the data was denoised by regressing out the top 1.5 % principal components of voxels that were outside of the brain. Additionally, the data was band-pass filtered between 0.005 and 0.5 Hz, eliminating slow drifts (>3 minutes) and fast fluctuations (<2 seconds), and a Savitzky–Golay filter was applied to temporally smoothen the data (2-second window). Finally, the fUS data was normalized to signal changes in percent by subtracting and dividing by the mean signal, yielding a relative measure of activity (ΔF/F or %) and, if indicated, z-scored.

### fUS behavior analysis

To generate voxel-wise activation maps shown in Fig. 2 around egress, grooming, and running, fUS data was first extracted in time windows surrounding each behavioral bout and averaged across all occurrences of a given behavior across sessions and mice. We report both first level and second level analysis (Extended Data Figures), but given the different number of animals for the two cohorts, we chose to report the one-level statistics in the main figure. For each behavior, the resulting activity was then averaged across time (0-10 seconds after egress onset, 0-15 seconds after grooming onset, 10-20 seconds after running onset) and tested for significance across all voxels using a one-sample t-test. When indicated in figure legends, p-values were corrected for multiple comparisons using the Benjamini-Hochberg procedure to control the false discovery rate (FDR). Thresholded significant voxels (thresholds reported in figure legends) were visualized on coronal slices of the Allen Brain Institute’s reference atlas. To determine general behavior switching nodes, we overlapped the top 20% significant voxels (cutoff p < 0.05) before transition to egress and running and plotted the shared voxels on top of the reference background. To examine the effect of behaviors on single-regions, the voxel-wise fUS data was segmented into regions. To this end, we collapsed all regions from the Allen Brain Institute’s mouse brain atlas into 134 brain regions and averaged voxel-data within each of these regions. These region-wise signals were visualized as heatmaps aligned to behavior onset. For further analysis, regions (ROIs) showing significant voxel-wise responses were selected, and their mean and standard error of the mean were plotted across time. To test whether fUS activity in these ROIs differed between baseline and the pre-transition period, average activity during a 5-second baseline 20 seconds before the onset was compared to activity during the 5 seconds preceding the onset of egress or running using a Wilcoxon signed-rank test.

### Whole-brain decoding

To determine whether there is a pattern in the brain that predicts transitions in behavior before their onset we have used the CanlabCore toolbox^42^ developed for application of SVMs on fMRI data. The code was adapted in a custom MATLAB script to accommodate fUS data. Following the approach used for decoding behavioral readouts, we extracted time-aligned voxel-wise fUS activity (111720 voxels) around transitions to egress, grooming, and running (120 frames). Moreover, 50 sets of pseudo-bouts were generated by shuffling behavioral onsets across sessions, preserving bout count and distribution while disrupting alignment with real behavior transitions. For each set of pseudo-bouts (50) and each time point (∼2 per sec), we trained a SVM classifier to distinguish real behavior bouts from pseudo-bouts based on voxel-wise activity patterns of the mouse brain. Each classifier was trained on 90% of the data and tested on the remaining 10%, yielding a 10-fold cross-validated accuracy score per time point. This process was repeated across all 50 pseudo-bout iterations, resulting in a distribution of decoding accuracies per time point. The average decoding accuracy and standard error of the mean were computed across these 50 models and plotted for all time points in Fig. 3. As a control, we decoded one set of pseudo-bouts against 49 different sets of pseudo-bouts per time point following the same procedure, and plotted the outcome as mean and standard error of the mean. Next, we calculated the EDT in the same way as for behavior decoding, taking the first time point, moving from behavior onset backward in time, at which the mean decoding accuracy overlapped with the control’s standard deviation. This allowed us to determine whether distributed voxel-level fUS activity predicted behavior prior to its onset, and to pinpoint when prediction accuracy first exceeded control levels. To assess whether the decrease observed via fUS in the MS, SSp, Sub and PAG might co-occur with the increase of predictability of behavior transitions from the whole-brain data, we calculated Spearman’s rank correlation between the fUS signals of these regions and the SVM decoding accuracy.

### Analysis of optogenetics data

To analyze the effect of optogenetic inhibition on whisking, pupil size, and VBA or wheel motion, we first extracted behavioral readouts from each session. We used Facemap^34^ to extract the pupil diameter and the motion SVD of the whisker region of each mouse. For extraction of wheel motion we used a MATLAB custom-code that computes the motion energy of a ROI of the wheel, allowing us to measure motion speed in one single trace per session. The VBA motion was analyzed directly from the VBA’s laser displacement sensor. Next, we pooled all trials of the VBA and wheel datasets, computed average motion and pupil traces (z-scored to the entire 60-second baseline), and plotted them in Fig. 4. For quantification, z-scored data was averaged during the stimulation period across animals. This data was tested for a significant difference between the control (mCherry) and opsin (stGtACR2) groups using a Wilcoxon rank-sum test for independent samples. Note that for the VBA data, a significant difference was only found when comparing the first 30 seconds after stimulation onset between the two groups. Grooming in the VBA was classified as described in the “Behavior classification” section above, by correlating a baseline ROI around the snout to every frame of the video and identifying frames in which the correlation coefficient dropped. In the running wheel dataset, all videos were manually scored for grooming, since the automatic detection code was not optimized for this setup. To this end, all videos were watched at 3x speed and grooming instances were marked. Next, we plotted the time spent grooming before and during stimulation across all trials that included grooming in Fig. 4, and quantified it using a Wilcoxon matched-pairs signed-rank test for both the control and opsin cohorts. For the running wheel dataset, we also tested for a difference in the number of grooming bouts occurring in the ON vs. OFF periods using a Wilcoxon matched-pairs signed-rank test. This was not done for the VBA dataset due to the smaller number of groom bouts, which was limited by the short session duration. Finally, we compared the duration of groom bouts occurring in the stimulation vs. non-stimulation periods using a Wilcoxon rank-sum test.

## Acknowledgements

We thank the lab members of the Macé lab, Oliver Barnstedt, and Nadine Gogolla for helpful discussions and feedback on the manuscript. We thank B. Apgaua and A. Ablitip for assistance with the analysis of the behavioral data, and N. Gogolla for providing us with viruses and resources for starting the project. We thank G. Montaldo and A. Urban for technical support with fUS data acquisition. We thank the MPI-BI and UMG animal facilities, veterinarians, and caretakers for their support. We thank the MPI-BI Imaging Facility and Histolab for assistance with histology and imaging. We thank Julia Kuhl for help with illustrations. B.J.E. was funded by the European Molecular Biology Organization Postdoctoral Fellowship no. ALTF 449-2020. E.M. was funded by the Max Planck Society, the Else Kröner Fresenius Foundation through the ElseKröner Fresenius Center for Optogenetic Therapies, the Niedersächsisches Ministerium für Wissenschaft und Kultur through zukunft.niedersachsen, and the Deutsche Forschungsgemeinschaft (DFG, German Research Foundation) through Germany’s Excellence Strategy EXC 2067/1-390729940, the Project number 537609931(FOR 5807, project 1) and the Collaborative Research Center 1690 (Project B05).

## Author Contributions

Experiments were designed by P.W. and E.M. fUS and optogenetic experiments were performed by P.W. L.B. provided technical assistance with optogenetic experiments and histology. Behavioral data were analyzed by P.W. and P.T.M. fUS data were analyzed by P.W., E.M., and B.J.E. B.J.E. developed the code for the fUS decoding, and helped design experiments and supervise P.W.

J.M.M.P. contributed code for preprocessing of fUS data. The manuscript was written by P.W. and E.M.

## Competing interests

The authors have no competing interests to report.

## Data and code availability

All data and code will be made available on public repositories upon publication.

## Notes

### Competing Interest Statement

The authors have declared no competing interest.

