## Supplementary Figures for "A transition-prone brain state precedes spontaneous behavioral switching"

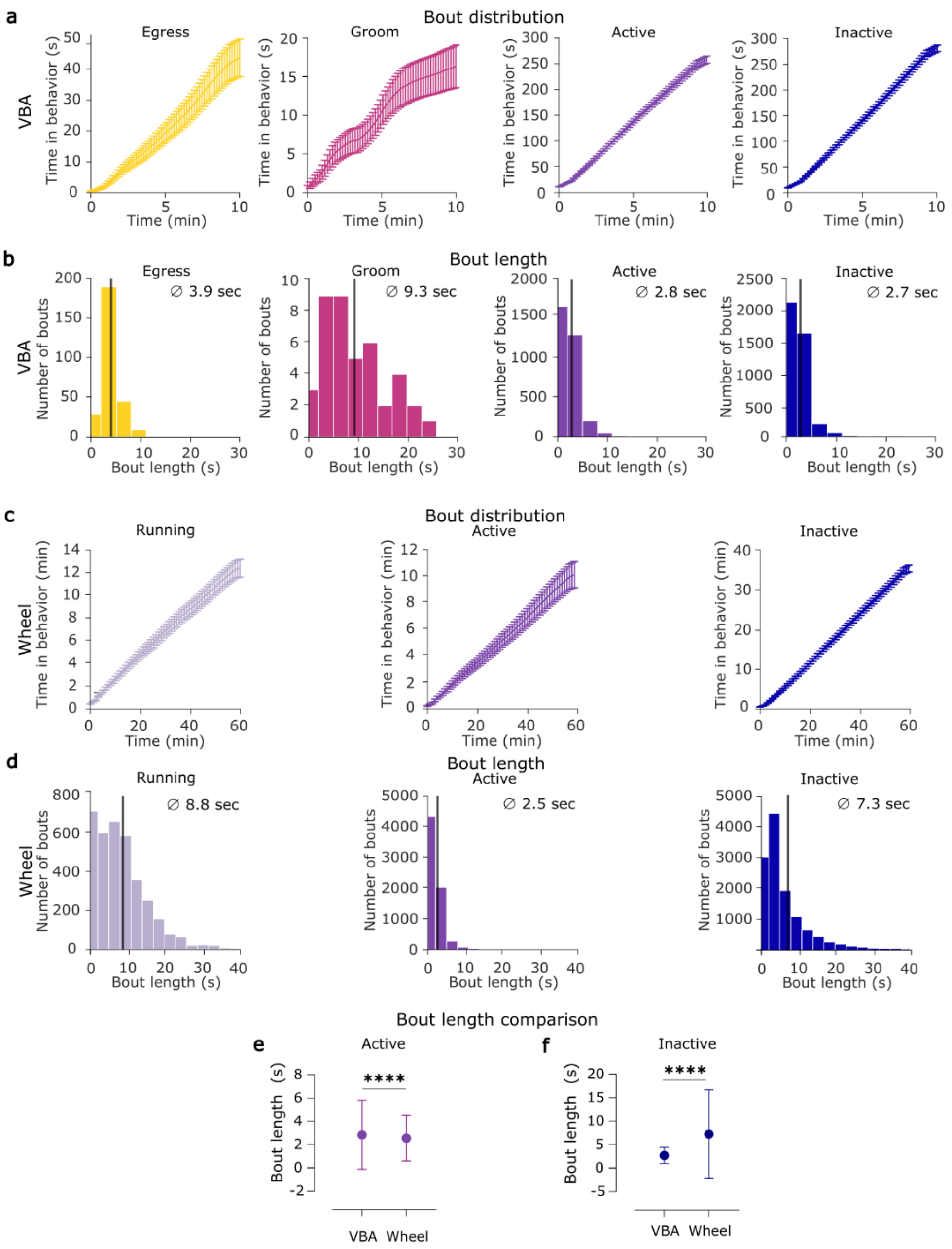


**Extended Data Fig. 1 | Bout distribution and bout length of behavioral states in the VBA and running wheel. a**, Cumulative sum of Egress, Grooming, Active and Inactive bouts. **b**, bout lengths of Egress, Grooming, Active and Inactive. **c**, Cumulative sum of Running, Active and Inactive bouts. **d**, bout lengths of Running, Active and Inactive. **e-f**, comparison of bout length of Active (e) and Inactive (f) bouts between bouts recorded in the VBA and in the running wheel setup. Mann-Whitney test. **** $p < 0.0001$.

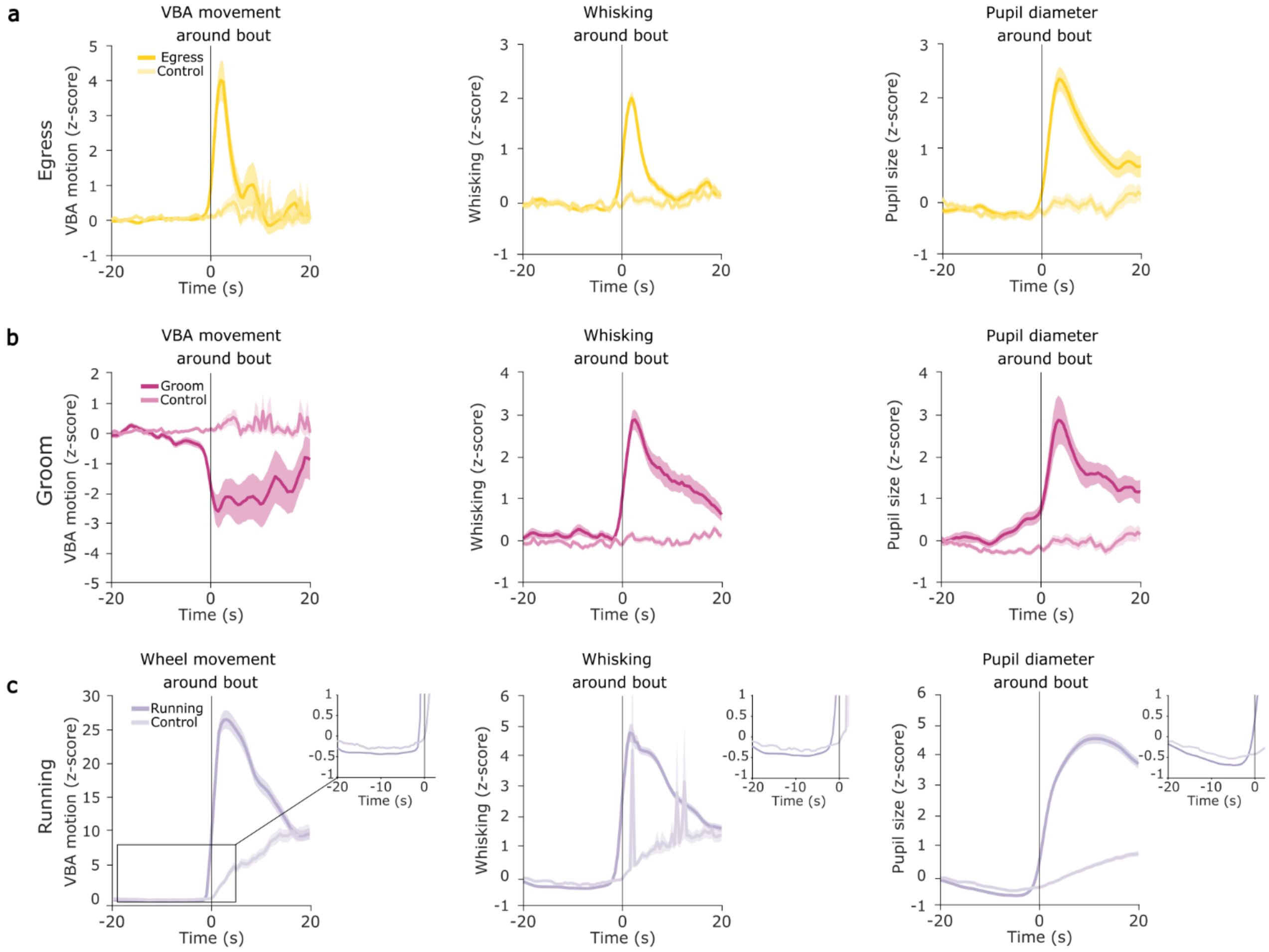


**Extended Data Fig. 2 | Behavioral readouts around behavior onsets with boutless baselines. a-c**, VBA motion (left), whisking (middle) and pupil diameter (right) around Egress bouts (**a**, n = 162 trials), Grooming (**b**, n = 44 trials), and Running (**c**, n = 1090 trials). Darker lines represent mean ± s.e.m. of behavior bouts and lighter lines represent pseudo-bouts as control here and elsewhere in the figure). The behavior starts at timepoint 0.

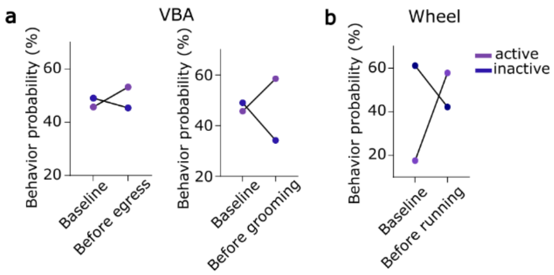


**Extended Data Fig. 3 | Probability of being Active before a behavior transition is increased. a**, averaged behavior probability in the baseline and one frame before egress (left) and grooming (right). **b**, average behavior probability around the onset of Running (left) and baseline vs one frame before Running comparison (right).

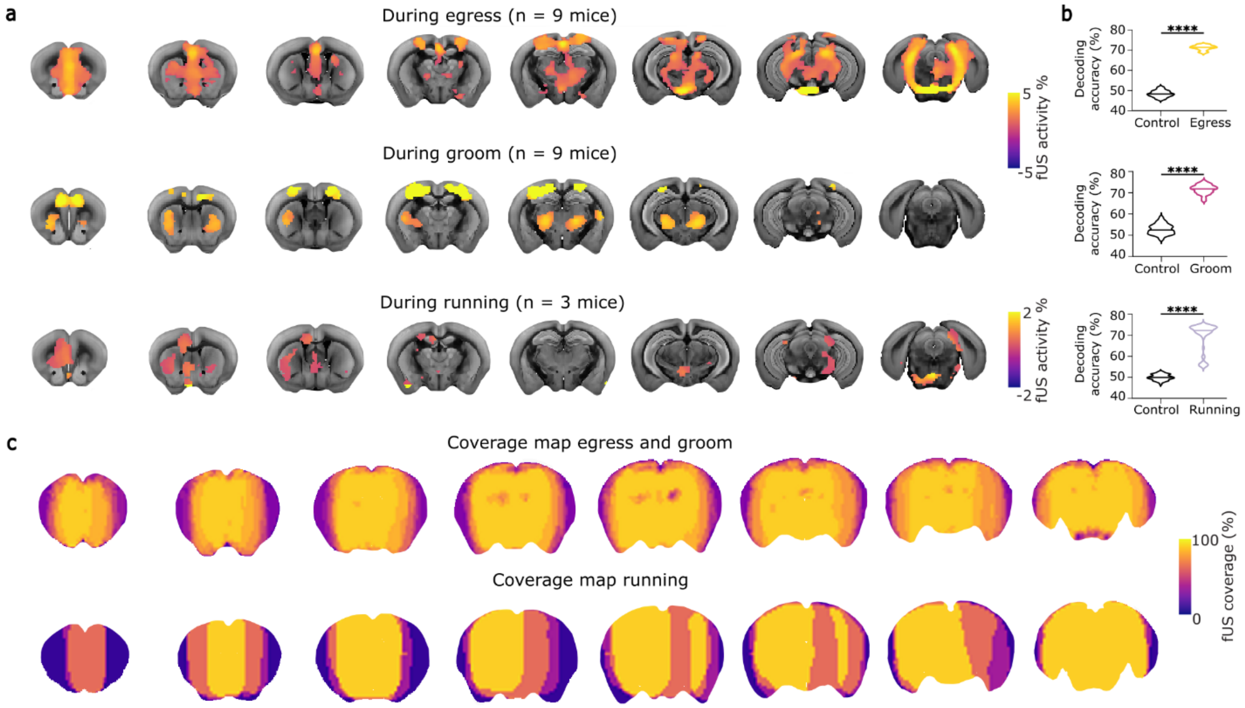


**Extended Data Fig. 4 | Whole-brain correlates of Egress, Grooming and Running (two-level analysis). a,** fUS activity patterns underlying Egress (**top**), Grooming (**middle**) and Running (**bottom**) displayed on reference coronal brain images. Voxels falling within threshold are shown: one-sample t-test, for Egress (n = 9 animals) FDR-corrected (FDR = 0.1) voxels, for Grooming (n = 9 animals) uncorrected ($p<0.02$), for Running (n = 3 animals) uncorrected ($p<0.1$). **b,** whole-brain decoding using SVM classification decoding behavior vs. pseudo-bouts for Egress (top), Grooming (middle) and Running (bottom) (50 SVM runs with 50 sets of pseudo-bouts, 10-fold cross-validation per run, n = 162 Egress bouts, n = 44 Groom bouts from 9 animals, and n = 399 bouts for Running from 3 animals here and elsewhere in the figure) tested against control (50 runs pseudo-bouts vs. pseudo-bouts), Wilcoxon paired sign rank test. **** $p < 0.0001$. **c**, fUS window coverage across all animals for Egress and Groom (**top**, n = 9 animals) and for running (**bottom**, n = 3 animals) in percent.

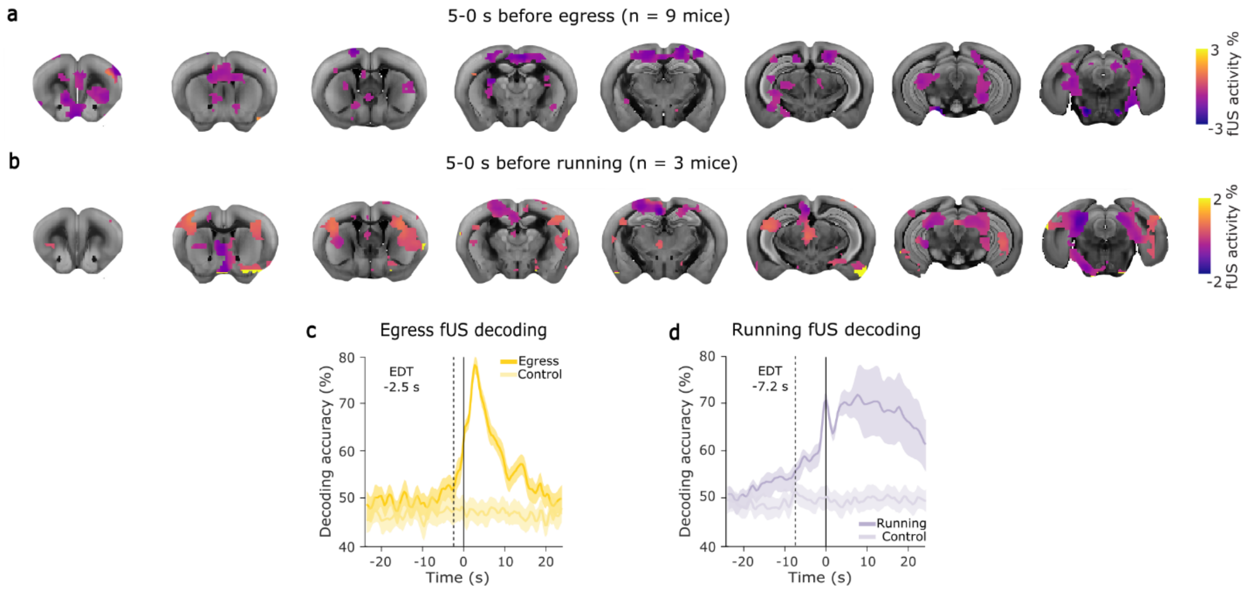


**Extended Data Fig. 5 | Whole-brain activity pattern before the onset of Grooming. a**, fUS activity patterns 5-0 seconds before the onset of Grooming (n = 44 bouts from 9 animals) on reference coronal brain images. Uncorrected, p < 0.1, one-sample t-test. **b**, fUS activity patterns 5-0 seconds before the onset of Grooming (n = 44 bouts from 9 animals) on reference coronal brain images. Uncorrected, p < 0.1, one-sample t-test. **c**, Whole-brain decoding (SVM classification behavior bouts vs. pseudo-bouts) using fUS data around Groom (n = 44 from 9 animals). Each timepoint represents an average decoding accuracy ± s.e.m. across 50 SVM runs each using a different set of pseudo-bouts. Control decoding accuracies (lighter pink) were generated by classifying one set of pseudo-bouts against 49 other sets of pseudo-bouts. Dashed line indicates the EDT, defined as the first time point (moving from during behavior to before) at which the mean decoding accuracy (behavior bout vs. pseudo-bout) falls within one standard deviation of the control decoding accuracy (pseudo-bout vs. pseudo-bout). The EDT for Groom is -3.5 seconds.

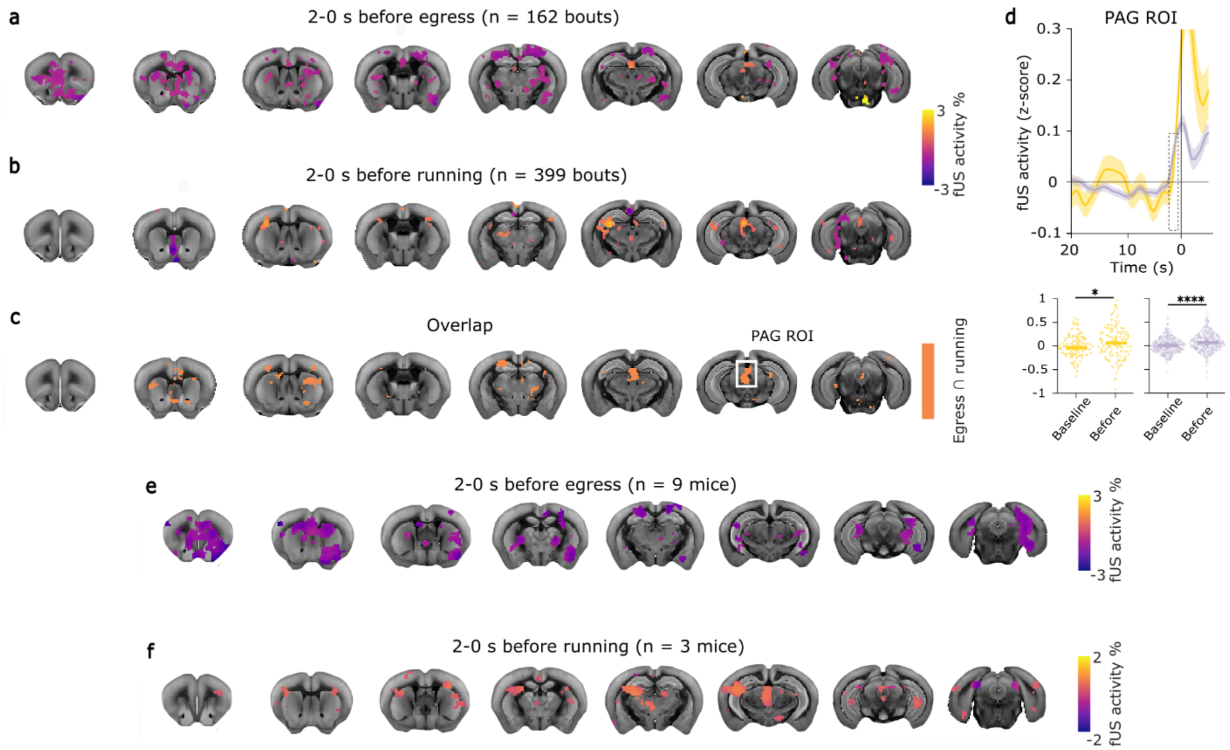


**Extended Data Fig. 6 | Whole-brain activity pattern 5 - 0 s before the onset of Egress and Running (two-level analysis). a-b**, fUS activity patterns 5-0 seconds before the onset of Egress (**a**, n = 9 animals) and Running (**b**, n = 3 animals) on reference coronal brain images. Uncorrected, p < 0.2 for Egress, p<0.3 for Running, one-sample t-test. **c-d**, Whole-brain decoding (SVM classification behavior bouts vs. pseudo-bouts) using fUS data around egress (c, n = 162 from 9 animals here and elsewhere in the figure) and running (d, n = 500 from 3 animals). Each timepoint represents an average decoding accuracy ± std. across 50 SVM runs each using a different set of pseudo-bouts. Control decoding accuracies (lighter yellow/gray) were generated by classifying one set of pseudo-bouts against 49 other sets of pseudo-bouts. Dashed lines indicate the EDT, defined as the first time point (moving from during behavior to before) at which the mean decoding accuracy (behavior bout vs. pseudo-bout) falls within two standard deviations of the control decoding accuracy (pseudo-bout vs. pseudo-bout). With this more stringent threshold the EDT for egress (c) is -2.5 seconds and for running (d) -7.2 seconds.

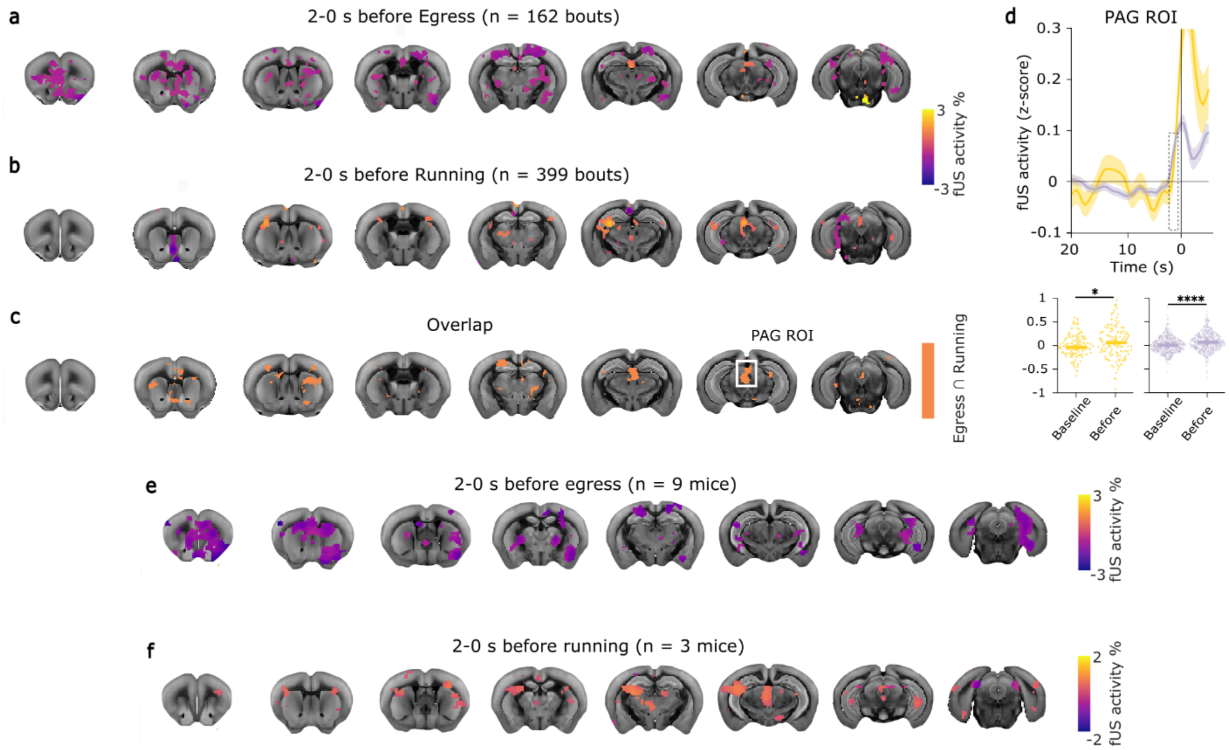


**Extended Data Fig. 7 | Whole-brain activity pattern 2 seconds before the onset of Egress and Running. a**, fUS activity patterns 2-0 seconds before the onset of Egress (n = 162 bouts from 9 animals) on reference coronal brain images. Only voxels significantly different from zero are shown, uncorrected, $p < 0.01$, one-sample t-test. **b**, fUS activity patterns 2-0 seconds before the onset of Running (n = 399 bouts from 3 animals) on reference coronal brain images. Only voxels significantly different from zero are shown, FDR-corrected (0.1), $p < 0.002$, one-sample t-test. **c**, Common significant regions 2 seconds before the onset of Egress and running. Overlap of 20% most significant voxels depicted in orange. White box highlights a common node: the PAG. **d**, fUS activity of ROI in PAG before onset of Egress (yellow line, mean ± s.e.m) and Running (gray line, mean ± s.e.m). Dashed box highlights 2-0 seconds before behavior onset. Bottom: fUS activity during baseline episode and before Egress (left, yellow) and Running (right, gray). Horizontal lines show median fUS activity. Wilcoxon signed-rank test. * $p < 0.05$, **** $p < 0.0001$. **e-f**, fUS activity patterns 2-0 seconds before the onset of Egress (**e**, n = 9 animals) and Running (**f**, n = 3 animals) on reference coronal brain images. Uncorrected, $p < 0.2$ for Egress, $p<0.3$ for Running, one-sample t-test.

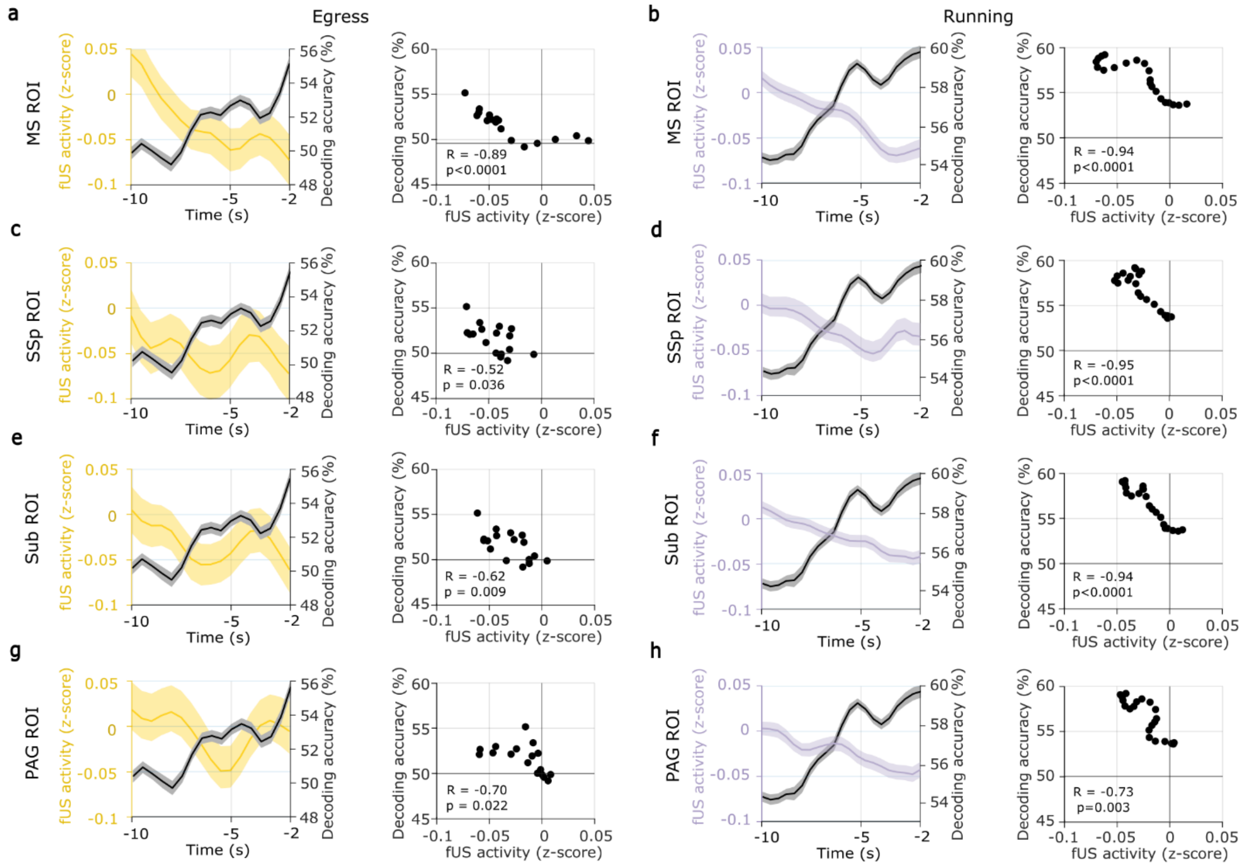


**Extended Data Fig. 8 | Decrease in fUS activity in selected regions is positively correlated with predictability 10 to 2 seconds before behavior onset**. **a-h**, fUS activity (yellow line, mean ± s.e.m. for Egress n = 162 bouts and gray line, mean ± s.e.m. for Running n = 399 bouts) of ROIs in MS (**a**,**b**), SSp, (**c**,**d**), Sub (**e**,**f**) and PAG (**g**,**h**) and whole-brain decoding accuracies across 50 SVM runs (black line mean ± s.e.m before behavior onset).

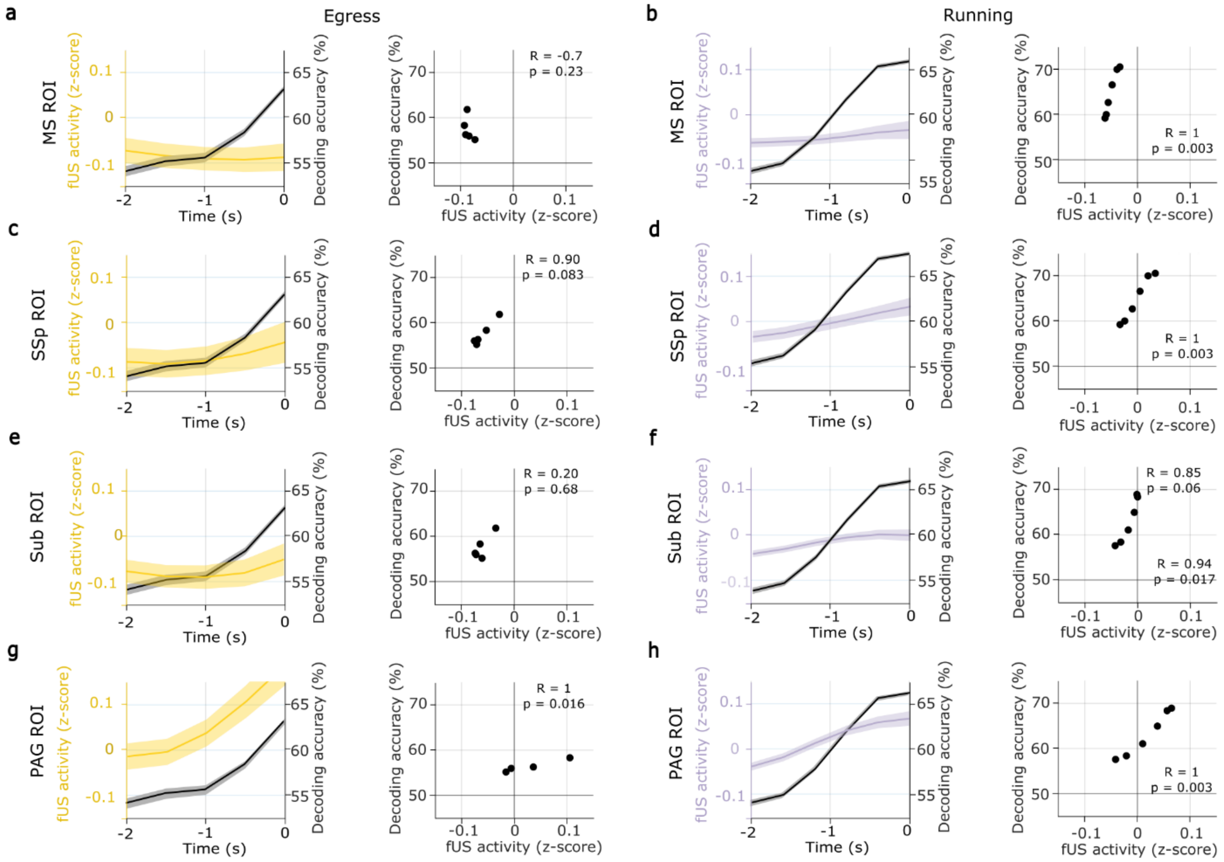


**Extended Data Fig. 9 | Decrease in fUS activity in selected regions is positively correlated with predictability 2 seconds before behavior onset**. **a-h**, fUS activity (yellow line, mean ± s.e.m. for Egress n = 162 bouts and gray line, mean ± s.e.m. for Running n = 399 bouts) of ROIs in MS (**a**,**b**), SSp, (**c**,**d**), Sub (**e**,**f**) and PAG (**g**,**h**) and whole-brain decoding accuracies across 50 SVM runs (black line mean ± s.e.m before behavior onset).

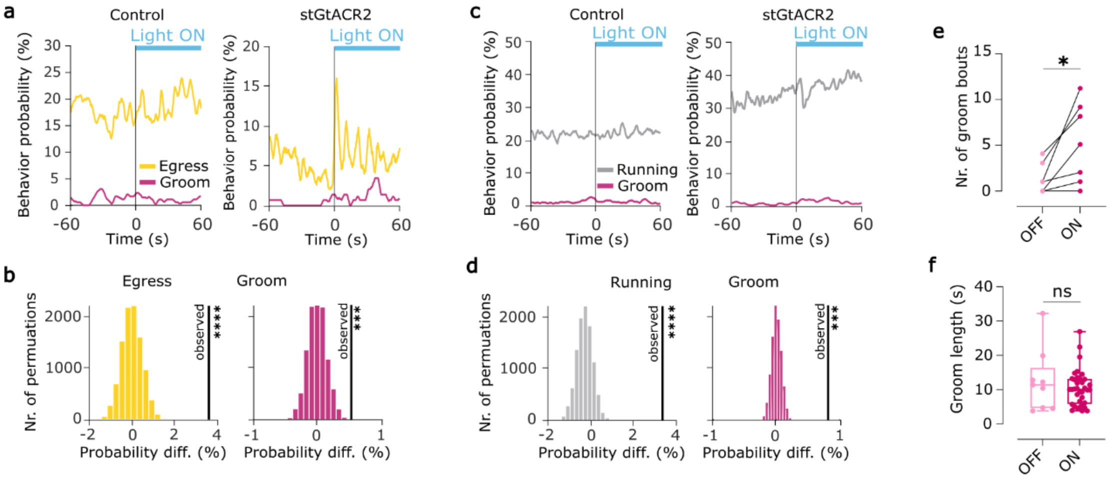


**Extended Data Fig. 10 | Probability of the states Egress, Grooming and Running is increased during optogenetic inhibition of MS. a, c,** Behavior probability of the behavior states Egress (yellow line), Grooming (pink line) and Running (gray line) in the control and stGtACR2 group before and during light stimulation in VBA (**a**) and on running wheel (**c**). **b**, **d**, Comparison of observed difference (vertical black line) in behavior probability of Egress and Grooming (**c**) and Running and Grooming (**d**) between baseline and light stimulation period with a control distribution created by shuffling behavior labels. **e**, Number of groom bouts during 60 second light stimulation period and baseline on the running wheel (n = 7 mice, Wilcoxon matched-pairs signed-rank test, $p = 0.031$). **f**, Grooming length in seconds during baseline and light stimulation period on the running wheel (Wilcoxon rank sum test, $p = 0.786$). * $p < 0.05$, *** $p < 0.001$, **** $p < 0.0001$.
